# Single cell snapshot analyses under proper representation reveal that epithelial-mesenchymal transition couples at G1 and G2/M

**DOI:** 10.1101/2025.02.24.639880

**Authors:** Sophia Hu, Yong Lu, Gaohan Yu, Zhiqian Zheng, Ke Ni, Amitava Giri, Jingyu Zhang, Yan Zhang, Guang Yao, Jianhua Xing

## Abstract

Numerous computational approaches have been developed to infer cell state transition trajectories from snapshot single-cell data. Most approaches first require projecting high-dimensional data onto a low-dimensional representation; however, this can distort the dynamics of the system. Using epithelial-to-mesenchymal transition (EMT) as a test system, we show that both biology-guided low-dimensional representations and trajectory simulations in high-dimensional state space, not representations obtained with *brute force* dimensionality-reduction methods, reveal two broad paths of TGF-β induced EMT. The paths arise from the coupling between cell cycle and EMT at either the G1 or G2/M phase, contributing to cell-cycle related EMT heterogeneity. Subsequent multi-plex immunostaining studies confirmed the multiple predicted paths at the protein level. The present study highlights the heterogeneity of EMT paths, emphasizes that caution should be taken when inferring transition dynamics from snapshot single-cell data in two- or three-dimensional representations, and shows that incorporating dynamical information can improve prediction accuracy.

## Introduction

Throughout the lifetime of every cell, there are critical moments when they encounter intrinsic and extrinsic signals that direct them towards a specific cellular fate. These decisions occur in various contexts, such as embryonic development, wound healing, cellular senescence, and aging. Within a cell multiple cellular processes function simultaneously; therefore, the decision and execution of a cell fate change often requires coordination between different cellular programs to allocate limited resources and to minimize interference or conflict.

A central, yet long-standing question in cell biology revolves around how the cell cycle couples to and influences other cell fate decisions such as differentiation, apoptosis, and reprogramming^1^. The cell cycle is a tightly regulated cellular process, where a proliferative cell cycles through a temporally ordered G1→S→G2→M sequence, with multiple checkpoints deciding on the continuation or halting of the cell cycle to ensure proper cell division.

The cell cycle couples with cell fate in a context-dependent manner. In the case of terminal differentiation, evidence has shown that cells typically withdraw from the cell cycle and enter an arrested state to proceed toward a specific cell fate^2^. In other cases, cell division is required to allow for cell fate switching, where the period between the end of mitosis and before S-phase entry is a prime window for transcriptional and epigenomic changes to lead to a change in cell fate. The Rb-E2F network, for example, integrates signals from regulators of other cellular processes to decide whether a newly divided cell proceeds through to the G1 phase or exits the cell cycle and enters a quiescent G0 phase^1,3^. In other cases, the specific cell cycle phase a cell resides in can dictate a cell’s fate^4–7^. For example, Gomer and Firtel found that single amoebae differentiated according to its position in the cell cycle, becoming prespore if in the G1 phase and prestalk cell if in the G2 phase^7^. The coupling between the cell cycle and cell fate changes has been exploited to modulate cell fate transitions, such as accelerating the reprogramming process and improving the efficacy of cancer treatment^6,8^. However, the exact mechanisms that govern cell fate decisions, specifically the timing along the cell cycle, remain elusive.

Epithelial-to-mesenchymal transition (EMT) is a cell phenotype transition that has attracted extensive attention in recent years, due to its involvement in many biological processes including development, wound healing, tissue fibrosis, and cancer metastasis^9^. During EMT, epithelial cells, characterized by an apical-basal polarity and adherence to a basement membrane and other epithelial cells, lose their epithelial traits and develop mesenchymal traits, characterized by a detachment from the basement membrane, an elongated shape and increased mobility^9,10^. The onset of EMT is regulated by a core regulatory network involving transcription factors and microRNAs. These regulators are also involved in other cellular processes like proliferation and regulation of the cell cycle^11–13^.

An open question surrounding EMT is whether it proceeds through a single path or multiple paths^9^. Pseudotime analyses of single cell RNA-sequencing (scRNA-seq) data of human mammary epithelial (MCF10A) cells treated with transforming growth factor-beta (TGF-β), to induce EMT, concluded that EMT proceeds through a one-dimensional continuum^14^. Markov transition model analyses on snapshot proteomic data of non-small-cell lung carcinoma cell lines treated with TGF-β, also reached a similar conclusion^15^. On the other hand, Wang et al. performed live-cell imaging studies of TGF-β induced EMT in a A549, human lung carcinoma epithelial cell line, with endogenous vimentin-RFP labeling in a multi-dimensional composite cell feature space^16,17^ and found at least two parallel paths that connect the initial epithelial state to the final mesenchymal state.

The cell cycle is known to couple with EMT and could therefore be a potential driver for the multiple paths. In the context of human oral squamous cell carcinoma cells TGF-β treatment induced G1 cell cycle arrest^18^. In the case of kidney fibrosis and pancreatic ductal adenocarcinoma, a link between cell cycle arrest and partial EMT has been established where *Snai1*-induced partial EMT also induces G2/M cell cycle arrest^19,20^. In other instances, such as during epidermal stem cell differentiation, there is a complete decoupling^21^. The discrepancy in the reports raises the question of whether the coupling between the cell cycle and EMT is context dependent. Another factor to consider is the strength of the coupling. In the case of a tight coupling, one would expect cells to arrest from very specific and narrow regions of the cell cycle, such as a checkpoint (Fig. 1a). In contrast, a loose coupling would result in a broad range of cells undergoing EMT in different stages of the cell cycle (Fig. 1b).

**Figure 1.**
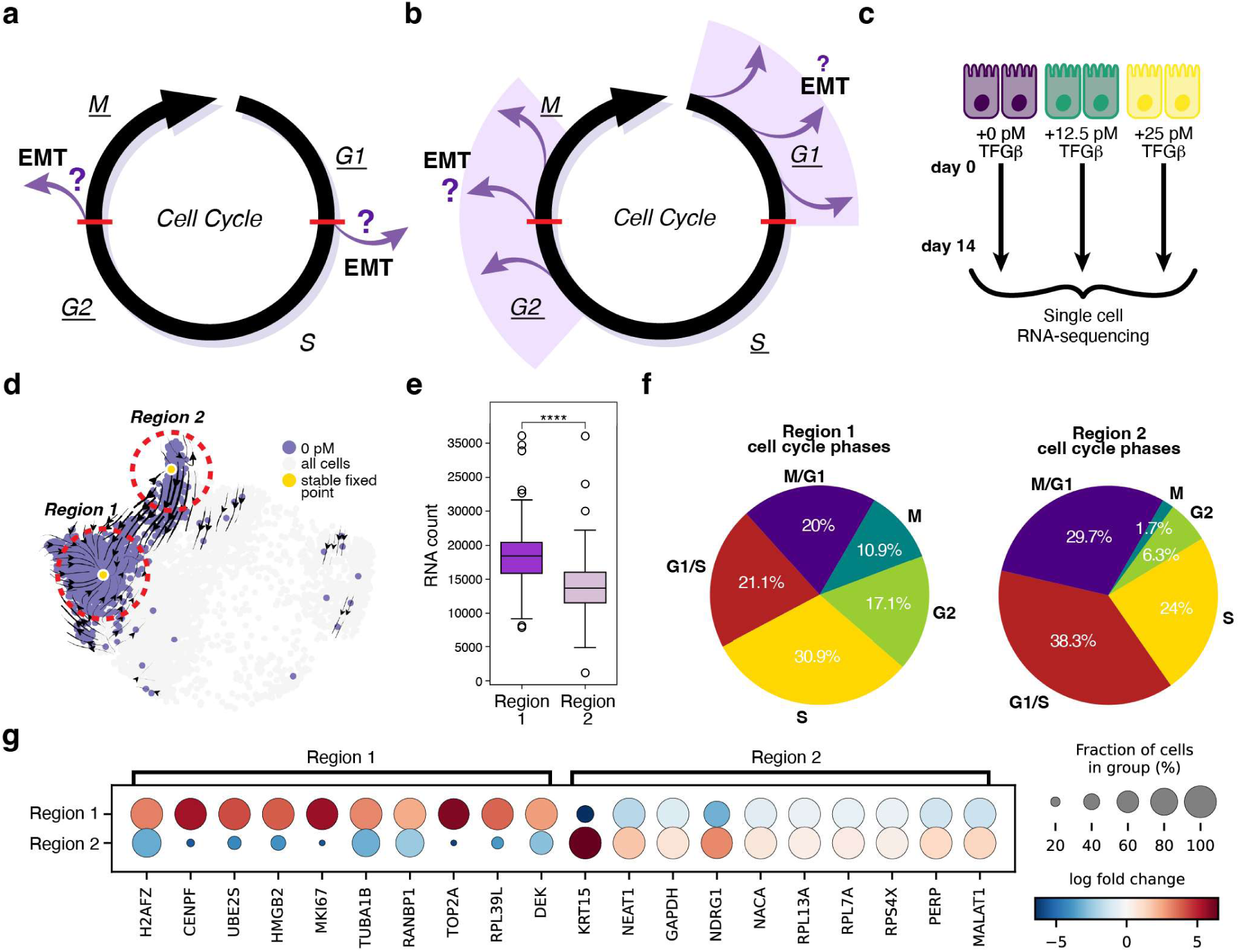
MCF10A cells undergo dose-dependent TGF-β induced EMT. (**a, b**) Plausible scheme of tight (**a**) and weak (**b**) coupling between cell cycle and EMT. Cell cycle checkpoints are indicated by the red lines. (**c**) Schematic of the experimental design of scRNA-seq data acquisition. MCF10A cells treated with increasing concentrations of TGF-β over the span of 14 days, pooled together and then scRNA-seq was performed. (**d**) Dynamo transcriptomic vector field of untreated MCF10A cells with stable fixed points. Circled regions are regions used for comparative analyses. (**e**) Comparison of the mRNA count between region 1 and region 2. (**f**) Comparison of the distribution of cell cycle phases between region 1 and region 2 using cell cycle phases assigned by dynamo. (**g**) Top ten differentially expressed genes between region 1 and region 2.

Addressing these questions is a formidable challenge as it requires unraveling the regulatory mechanisms and transition dynamics of cell phenotypic transition processes which involve many molecular species that are coupled together. The development of scRNA-seq technology has revolutionized biology research by providing genome-wide snapshots of gene expression profiles in heterogeneous cell populations^22,23^. Numerous computational approaches have been developed to infer phenotype transition trajectories from snapshot single-cell data. Pseudotime analysis is one class of computational approaches, where sampled single-cell data are ordered based on their progression through a biological process determined by gene expression similarities^24^. Methods such as optimal transport use distributions in the cell state space measured at several time points to infer how cells transition between different cell states^25^.

Another class of computational methods integrates single-cell expression states with dynamical information for trajectory inference. In a seminal study, La Manno et al. took advantage of the ability to distinguish unspliced and spliced mRNA transcripts from scRNA-seq data, and estimated instantaneous RNA velocity, the speed and direction of change of a high-dimensional vector in the cell expression space^26^. By taking these instantaneous RNA velocities, estimated using the original splicing-based method or by other improved methods^27^ as input, Qiu et al. developed *dynamo*, a machine learning framework that reconstructs a set of multi-dimensional, analytical, and generally nonlinear continuous vector field functions encoding the genome-wide gene regulation relations of the system^28^. Zhang et al. further developed *graph-dynamo* for complete dynamical model studies^29^.

In general, many of the existing dynamics-inference methods are performed on an extremely low-dimensional (e.g., two- or three-dimensional) manifold of the single-cell data^24^. That is, one implicitly assumes that the very low-dimensional representation faithfully distinguishes different cell states in a biologically meaningful manner. However, this assumption needs to be carefully evaluated, as inappropriate representations may lead to a distortion of the underlying cellular dynamics^30,31^ and possibly imprecise conclusions which we will discuss later.

While preliminary analysis in low dimensional space can aid in visually understanding scRNA-seq data, more emphasis needs to be placed on performing higher dimensional analysis to corroborate conclusions made from low dimensional analysis. Similar to molecular dynamics simulations which use a force field to describe molecular interactions, with *dynamo* one can perform deterministic or stochastic simulations with the learned vector fields to generate single-cell trajectories in a high-dimensional (e.g., > 20) cell state space to better understand the dynamics within the data.

To better understand EMT dynamics, here we performed vector field analyses using *dynamo* on scRNA-seq data of MCF10A cells treated with increasing concentrations of TGF-β and analyzed the dynamic cellular paths. To explore the coupling between the cell cycle and EMT, we extended upon the cell cycle framework from the R package Revelio^32^ to construct a computational pipeline to obtain a cell cycle-informed representation. With this representation we predicted a loose coupling between the two processes exhibited by broad transition regions, originating from either the G1 or G2/M cell cycle phase. Next, we performed trajectory simulations and transition path analyses to confirm these paths in high-dimensional space. Finally, to validate these findings we performed iterative indirect immunofluorescence imaging (4i) measurements and confirmed the two predicted EMT paths. Overall, this study advances our knowledge on the paths cells can take during EMT by identifying two broad EMT paths characterized by the coupling between the cell cycle and EMT. These findings highlight the importance of not only obtaining biologically informed low-dimensional space representations but also performing analyses in high-dimensional space to accurately capture dynamics within scRNA-seq data.

## Results

### scRNA-seq data of TGF-β induced EMT in MCF10A cells revealed an EMT continuum

We first analyzed the scRNA-seq dataset of human mammary epithelial (MCF10A) cells generated by Panchy et al^33^. The MCF10A cells were treated with increasing concentrations of TGF-β, an inducer of EMT, (0, 12.5, 25, 50, 100, 200, 400, and 800 pM, respectively) for 14 days and then sequenced, encompassing a total of 8983 cells. To confirm EMT progression, we calculated an EMT score for each cell across the entire range of TGF-β concentrations using a Kolmogorov-Smirnov (KS) EMT scoring method, where a negative score indicates a cell as more epithelial while a more positive score indicates a cell as more mesenchymal^34^. As the TGF-β concentration increased the overall KS EMT score increased indicating that cells overall gained mesenchymal characteristics and lost epithelial characteristics (Fig. S1a), consistent with what was observed by Panchy et al.^33^

To visualize EMT progression in the gene expression space, we examined the single-cell data with a two-dimensional Uniform Manifold Approximation and Projection (UMAP) representation^35^ (Fig. S1b). With increasing TGF-β concentration the cells shifted from left to right, suggesting that the first UMAP mode (UMAP1) mainly reflects the TGF-β response, while the second mode (UMAP2) mostly captures TGF-β independent variations. Curiously, the four highest doses (100, 200, 400, 800 pM of TGF-β) settled in the same region on the 2D UMAP (Fig. S1b).

Observing the cells in 3D revealed that the cells with each increasing TGF-β concentration simultaneously moved upward along the UMAP3 axis as well as to the right along UMAP1 (Fig. S1c, d). Amongst the cells treated with highest TGF-β concentrations, both the KS EMT score and expression of *FN1*, a mesenchymal gene marker, varied within each TGF-β concentration reflecting the coexistence of a heterogeneous continuum of mesenchymal states instead of a singular one (Fig. S1e, f), consistent with what has been reported in other studies^33,36^. Overall, the 3D UMAP provides a clearer representation of EMT progression and reveals how the overall perception can be obscured by the distortion that can occur in 2D representations. This observation resonates with recent debates on the common practice of analyzing single-cell data in a reduced two-dimensional space^30,37^.

Among the eight TGF-β concentrations, only the cells treated with 12.5 and 25 pM of TGF-β retained a subpopulation of epithelial cells indicated by the overlap with the untreated cells, which only contain epithelial cells (Fig. S1g-i). The remaining TGF-β concentrations contained little to no epithelial cells as most cells have already undergone EMT and have already moved away from the epithelial region, occupied by the untreated cells. In subsequent analyses on understanding how cells transition from the epithelial to mesenchymal region, we focused on the two lowest TGF-β concentrations (12.5 pM and 25 pM) in addition to the untreated cells (0 pM) (Fig. 1c). Consistent with the analysis of all TGF-β concentrations, the cells treated with low concentrations of TGF-β were also undergoing EMT (Fig. S2a-f). However, there was less heterogeneity in the end mesenchymal state as most cells end up in a high KS EMT score and high *FN1* expression state (Fig. S2b, c). These results suggests that at lower concentrations of TGF-β cells reach a single mesenchymal state^38^.

### Two-dimensional transcriptomic vector field of untreated MCF10A cells revealed both quiescent and cycling cells but was biologically distortive

To analyze the EMT process, we began with the untreated MCF10A cells. We first applied *dynamo* to obtain the corresponding RNA vector field (Fig. 1d). The reconstructed vector field identified two stable fixed points, towards which vectors converged, representing stable cell states. To characterize the two stable cell states, we selected cells surrounding each fixed point and performed comparative analyses of mRNA abundance, cell cycle phase distribution and gene expression. Interestingly, cells in region 1 had significantly higher raw mRNA counts than cells in region 2 (with an independent sample two-sided T-test *p* value 9e^-19^) (Fig. 1e), indicative of higher total RNA content, although the correlation can be confounded by technical variations. Furthermore, cell cycle phase estimation analysis revealed that region 1 had 17.1 percent more cells in the G1/S phase and 9.7 percent more cells in the M/G1 phase than in region 2 (Fig. 1f). In addition, differential gene expression analysis revealed that region 1 had a positive log fold change for cycling related genes including *MKI67*, *TOP2A*, and *CENPF* (Fig. 1g). Collectively, these results suggest that region 1 corresponds to cells actively proliferating while region 2 may correspond to cells enriched in the quiescent (i.e., G0) state.

Although the 2D UMAP reduced representation of the untreated cells was able to identify two subpopulations of cells, quiescent and proliferating, the representation distorts the cell cycle dynamics. Specifically, the region 1 stable fixed point, identified on the 2D space is both biologically and mathematically unreasonable, as it represents an actively proliferating cell state. Instead, one would expect to observe circular vector flows to represent the proliferating states. To be more specific, with proliferative cells one would expect that the single-cell data form a closed geometric manifold with a hollow center along the axis of the closed structure, representing cell cycle progression in the cell state space as shown by Schwabe et al^32^.

### Two-dimensional transcriptomic vector field revealed two possible EMT transition regions in TGF-β induced EMT of MCF10A cells

A fundamental question in the field of EMT is whether cells undergoing EMT follow a single path or multiple paths^9^. To investigate this question, we first applied the pseudotime method, Monocle 3^39^, to the MCF10A cells treated with 12.5 or 25 pM of TGF-β. This analysis identified a single EMT path (Fig. 2a, b), consistent with previous reports using pseudotime analyses^14^ and Markovian transition model analyses^15^ on snapshot single-cell data but differs from reports from live-cell imaging studies^16,17^.

**Figure 2.**
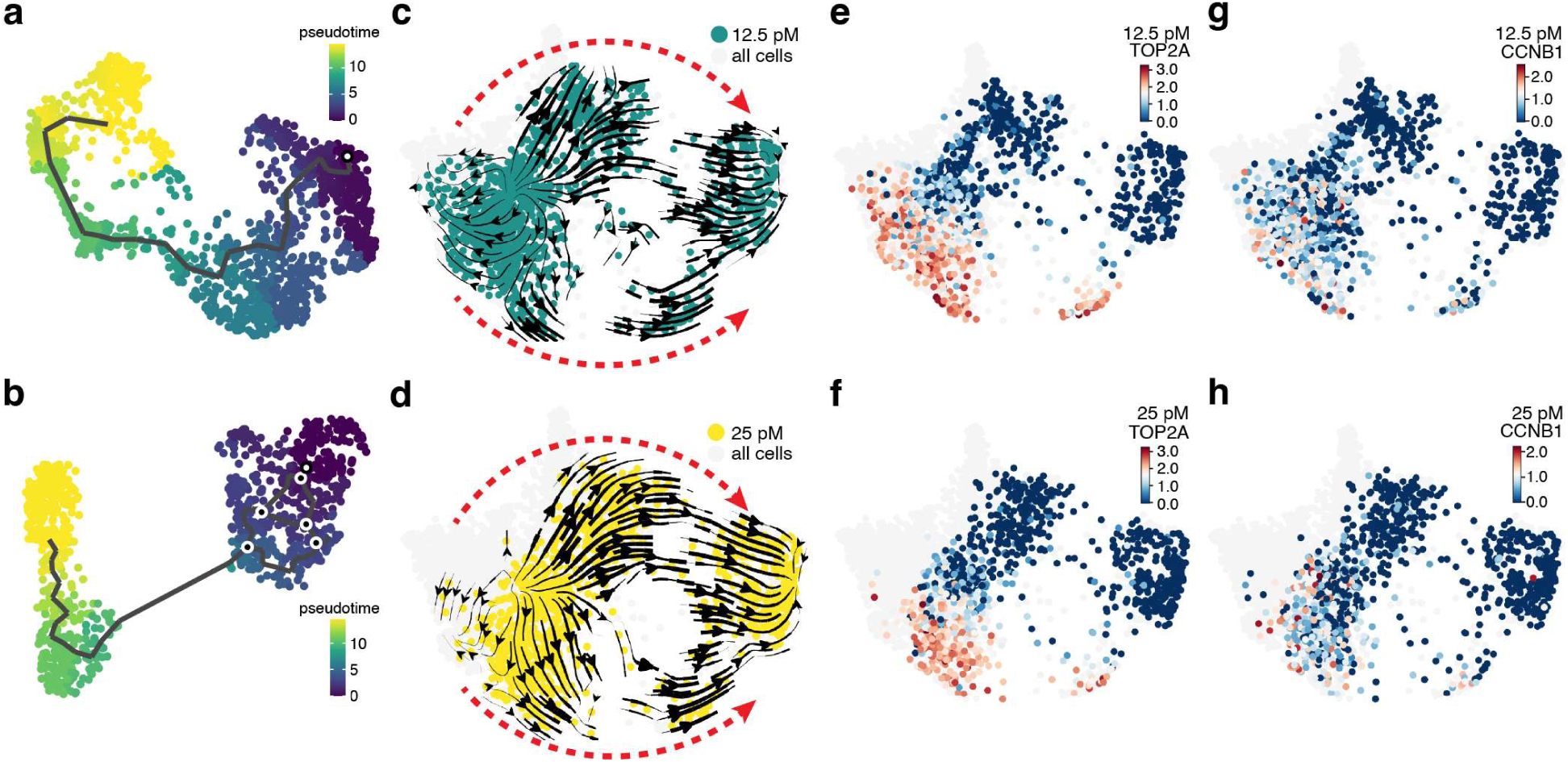
Vector fields of TGF-β induced MCF10A cells reveal two EMT paths. (**a, b**) Pseudotime analysis of 12.5 (**a**) and 25 (**b**) pM TGF-β treated cells on UMAP space. (**c**-**d**) Vector fields of 12.5 (**c**) and 25 (**d**) pM TGF-β treated cells on UMAP space. The red arrows indicate the two EMT paths. (**e**-**h**) TOP2A and CCNB1 gene expression for both the 12.5 (**e**) and 25 (**h**) pM TGF-β treated cells on UMAP space, respectively.

To investigate this discrepancy further, we performed vector field analysis using *dynamo*^28^. Notably, each vector field demonstrated a consistent pattern of EMT progression, with vectors shifting from left to right through two visually distinct transition paths (Fig. 2c, d), consistent with live-cell imaging results^16,17^. We noticed that several G2 cell cycle phase related genes, *TOP2A* and *CCNB1,* exhibited higher expression in the lower group of cells across the two concentrations of TGF-β treated cells (Fig. 2e-h). This observation led us to hypothesize that the upper and lower EMT paths observed from the vector fields were due to the cell cycle coupling with EMT at the G1/S and G2/M checkpoints (Fig. 1a, b).

However, the relationship between the cell cycle and EMT is unclear in the current two-dimensional UMAP representation (Box 1). Thus, one needs a representation that can simultaneously reflect cell cycle progression as well as EMT status of individual cells.

### Computational pipeline identified the Cell Cycle Coordinate

To determine whether the cell cycle and EMT couple together we required a representation that takes cell-cycle information into account, which is not explicitly accounted for in widely used dimensionality reduction methods. Therefore, we developed a four-step pipeline that builds upon the Revelio cell cycle framework from Schwabe et al^32^ to generate a biologically meaningful axis to capture cell cycle progression (Fig. 3a).

**Figure 3.**
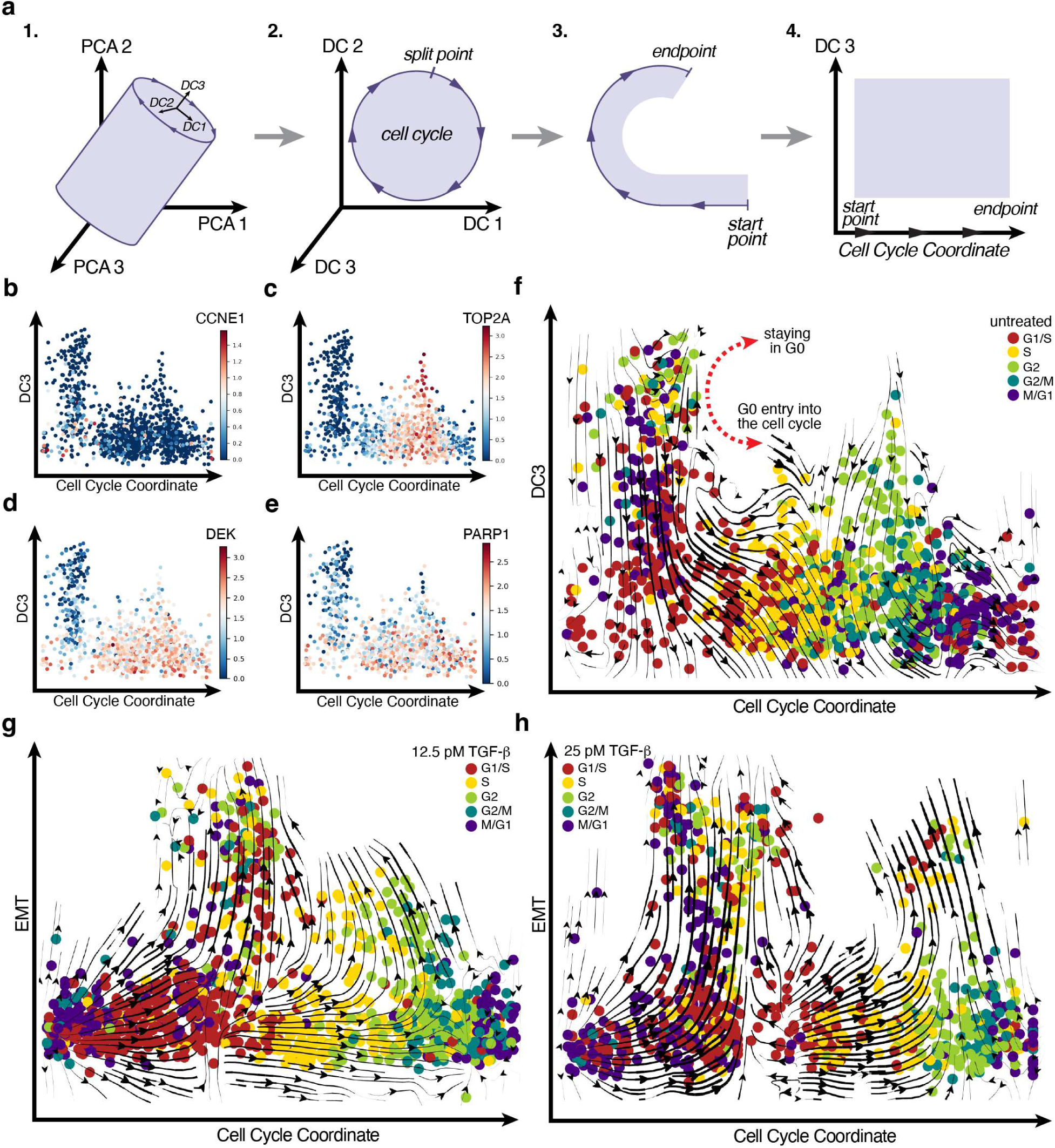
The cell cycle-EMT representation reveals broad cell cycle arrest regions. (**a**) The computational pipeline to obtain the Cell Cycle Coordinate. (**b**, **c**) CCNE1 (**b**) and TOP2A (**c**) gene expression to confirm cell cycle progression along the x-axis. (**d**, **e**) DEK (**d**) and PARP1 (**e**) gene expression confirming quiescence entry into the cell cycle along the y-axis. (**f**) The cell cycle-quiescent representation obtained from the untreated MCF10A cells reveals quiescent cell entry into the cell cycle. (**g**, **h**) The cell cycle-EMT representation obtained from the 12.5 (**g**) and 25 pM (**h**) TGF-β treated MCF10A cells. Colors of the data points in panels f-h represent discrete cell cycle stages were assigned using Revelio, which were shown to compare with the cell cycle coordinate.

The pipeline utilizes raw spliced scRNA-seq mRNA counts as input to Revelio^32^, an R package (Fig. 3a subpanel 1). Revelio assumes that the cell states of proliferative cells approximately reside on a hollow hypercylinder manifold. In step one, Revelio represents the data in a high-dimensional principal component (PC) space and then rotates the coordinate frame to define new axes referred to as dynamical components (DC). The first two DCs, DC1 and DC2 respectively, of the high-dimensional DC space encapsulate most of the cell cycle-dependent variation. In step two, we used an iterative finite temperature string method^16,40,41^ to obtain a one-dimensional cyclic cell cycle (CC) coordinate on the DC1-DC2 plane (Fig. 3a subpanel 2, Fig. S3a-f). Next, we identified the cell cycle division point, the split point, where we then unraveled the cell cycle coordinate to form a linear axis with a periodic boundary condition reflecting cell cycle progression (Fig. 3a subpanel 3 and 4). The remaining DCs orthogonal to the DC1-DC2 plane contain variations in the data that are mostly cell cycle independent. Thus, in our subsequent analyses, we employed the CC coordinate and the third dynamical component, DC3, to form a low-dimensional representation describing cell cycle progression and other cellular processes within the data (Fig. 3a subpanel 4). Details are provided in the method section.

### The cell cycle representation revealed cycling and quiescent dynamics

We first applied this computational pipeline to the untreated MCF10A cells. Indeed, single-cell data on the DC1-DC2 plane captured the cyclical nature of the cell cycle, which was further reflected in the vector field (Fig. S3g). The obtained CC coordinate accurately captured cell cycle progression, as the gene expression profiles of *CCNE1, TOP2A* and other cell cycle related genes showed expected elevated expression at either the G1 or G2/M phase of the CC coordinate (Fig. 3b, c, Fig S4a-c).

In addition, the cycling region previously identified in the UMAP space (region 1, Fig. 1d) was distributed across all cell cycle phases of the cell cycle when mapped to the new CC representation (Fig. S4d, e). The previously identified quiescent region, region 2 (Fig. 1d) ended up concentrated in the upper region of the DC3 axis in the new representation, where further down the DC3 axis marked the transition from quiescence to entry into the cell cycle (Fig. S4d, e). Furthermore, the gene expression of *DEK* and *PARP1*, genes identified as universally down-regulated during quiescence^42^, had reduced expression in the identified quiescent region and higher expression in the actively proliferating region (Fig. 3d, e).

The vector field obtained from the untreated MCF10A cells using the CC-DC3 representation further provided a dynamic view of cell cycle progression revealing a bifurcation between proliferation and quiescence (Fig. 3f). Above the bifurcation, the vector field flowed upward indicating cells progressing deeper into quiescence, while below the bifurcation, the downward vectors signal entry into the cell cycle. Such a bifurcation could reflect the quiescence-proliferation transition controlled by a *RB-E2F* bistable switch where after mitosis daughter cells either enter quiescence or proceed to the G1 phase of the cell cycle^43^. In conclusion, the CC-DC3 representation correctly captured the quiescence-proliferation dynamics.

### The cell cycle-EMT representation revealed coupling between the cell cycle and EMT

Next, we applied our computational pipeline to the 12.5 and 25 pM TGF-β treated MCF10A cells. Consistent with what was observed with the untreated cell data, the CC coordinate accurately depicted cell cycle progression, corroborated by high expression of *CCNE1* and *TOP2A* in the G1 and G2/M phase respectively (Fig. S5a-d). Gene expression analysis confirmed that the new y-axis effectively captures EMT progression, with decreasing gene expression levels of *CDH1*, an epithelial related gene (Fig. S5e, f) and increasing gene expression levels of *VIM* and *FN1,* mesenchymal related genes (Fig. S5g-j). That is, as cells moved upward along the EMT axis, they gradually lost their epithelial features and gained mesenchymal features.

Unlike the vector fields in the 2D UMAP representation for 12.5 and 25 pM TGF-β treated cells, the vector field on the CC-EMT representation depicted cells undergoing EMT originating broadly from either the G1 or G2/M phase (Fig. 3g, h, Fig. S5k, l). The widespread distribution suggested that the cell cycle and EMT were loosely coupled. If they were tightly coupled one would expect cells to exit from a narrow region of the cell cycle, likely corresponding to a cell cycle checkpoint.

Overall, our computational pipeline obtained a representation that captured cell cycle progression along a single axis, which enabled the extraction of valuable biological insight, specifically the broad coupling between the cell cycle and EMT.

### Trajectory simulations revealed two EMT paths originating from either G1 or G2/M in a multi-dimensional state space

To confirm that the coupling between the cell cycle and EMT at the G1 and G2/M cell cycle phases was not an artifact of dimensionality reduction, we performed trajectory simulations and transition path analyses in a 30-dimensional PCA space, where the vector fields were originally constructed. To obtain trajectories in high-dimensional space, we first identified an epithelial and mesenchymal region from the CC-EMT representation, designating them as potential start and end points, respectively (Fig. 4a). Using the transition matrix derived from the scRNA-seq data we launched trajectories from the epithelial start region and recorded trajectories that hit the mesenchymal end region, referred to as reactive trajectories.

**Figure 4.**
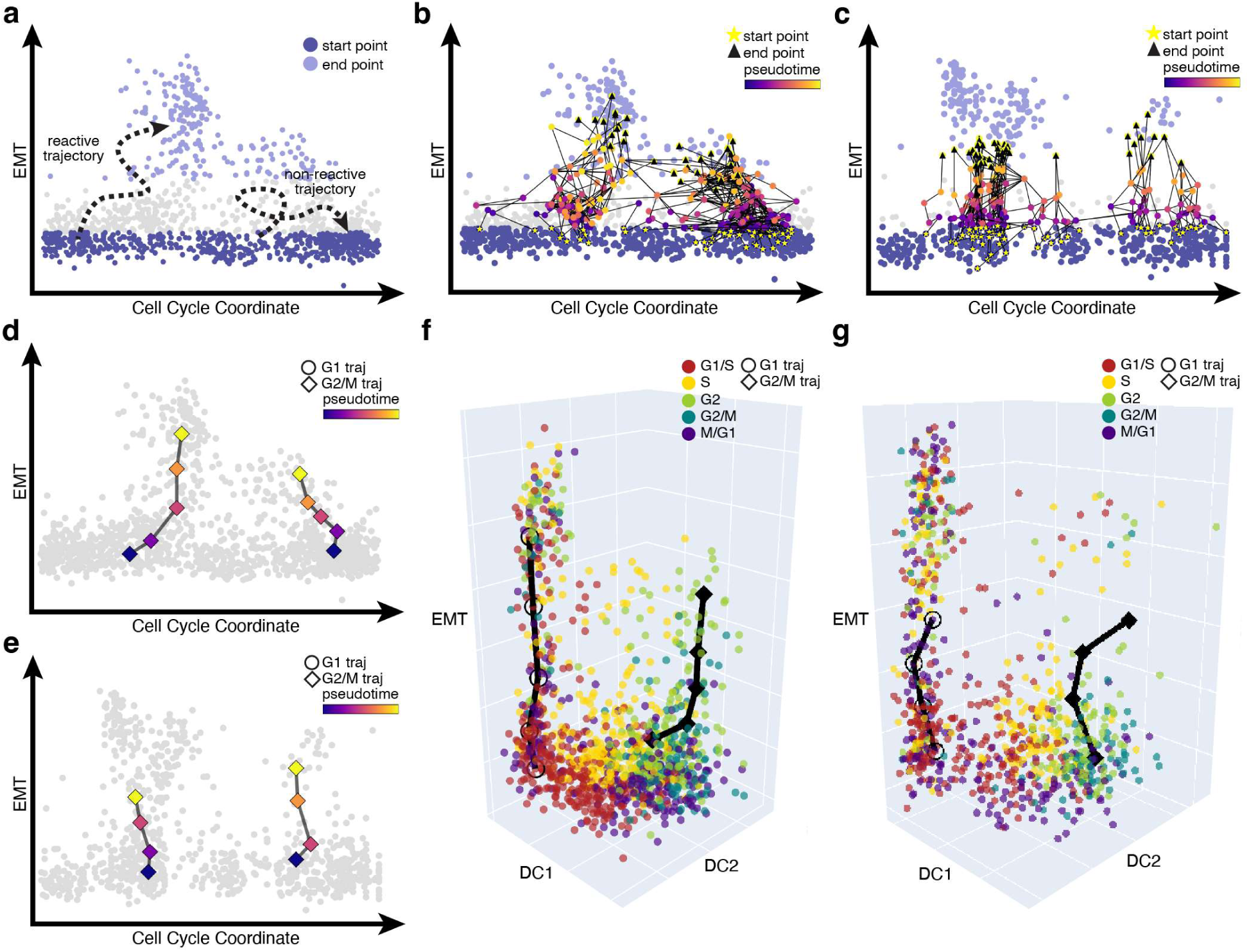
Analyses of trajectories simulated in 30-dimensional PCA space reveal two classes of EMT trajectories. (**a**) Trajectory simulation set-up for 12.5 pM TGF-β treated MCF10A cells with specified start (Epithelial) and end (Mesenchymal) region. (**b**, **c**) Example trajectories from one round of 12.5 pM (**b**) and 25 pM (**c**) TGF-β simulations mapped onto the EMT-Cell Cycle coordinate representation. (**d**, **e**) Mean trajectories from the (**d**) 12.5 pM and (e) 25 pM TGF-β simulations mapped on to the EMT-Cell Cycle coordinate representation. (**f**, **g**) Mean trajectories for two clusters in the 3-d DC-EMT space for the 12.5 pM (**f**) and 25 pM (**g**) TGF-β simulations. Cell cycle phases were assigned using Revelio.

For both the 12.5 and 25 pM TGF-β treated cells, we conducted ten rounds of trajectory simulations, generating 100 trajectories per simulation (Fig. 4b, c) and found that they grouped into two distinct clusters (Fig. S6a, b). With these two clusters of trajectories, we used a modified finite temperature string method^16,40^ to obtain the mean trajectory, similar to that of reaction coordinates in chemical reaction dynamics, per cluster. Projecting this ensemble of mean trajectories to the CC–EMT representation, we observed that the two clusters originated from either the G1 or G2/M phase (Fig. 4d, e). Inspection in 3D further confirmed that the two mean trajectories originated from either the G1 or G2/M phase (Fig. 4f, g).

It is important to note that the number of clusters was user defined, originally set as two, for both the 12.5 and 25 pM TGF-β trajectory simulations. We also tested whether the trajectories could be clustered into three groups. However, for both TGF-β doses the third mean trajectory aligned with either the already established G1 or G2/M trajectories, rather than forming a new path (Fig. S6c-f).

EMT-related genes between the G1 and G2/M paths both exhibited an increase in expression along the EMT axis. However, the G2/M path generally displayed lower expression at the beginning compared to the G1 path (Fig. S7). Additionally, *SNAI2,* an EMT transcription factor, *IL1B, S100A14, C1R,* all related to innate immunity, and *TNC*, which encodes a large extracellular matrix protein, showed differential expression between the two paths (Fig. S7). Further analysis is required to investigate the biological implications of these EMT path differences.

Overall, the trajectory simulation analyses confirmed that MCF10A cells undergoing TGF-β induced EMT could take one of two paths, transitioning broadly from either the G1 or G2/M region in high dimensional space.

### Multiplex immunostaining confirmed two transition regions in TGF-β treated MCF10A cells

Analyses of scRNA-seq from both 12.5 and 25 pM TGF-β treated MCF10A cells identified the G1 and G2/M phases as key regions where cells undergo EMT. To test whether these transition regions could be identified on the protein level, we treated MCF10A cells with 0.55 ng/ml (22.5 pM) of TGF-β for 0, 3, and 14 days (Fig. 5a). Cell morphology shifted from the typical round epithelial shape to an elongated, spindle-like form (Fig. 5b). These changes were classical indicators of EMT, confirming that EMT had occurred by 14 days of TGF-β treatment^38^. Not only did we observe a change in morphology, but we also observed changes in cellular markers. To characterize the changes in cellular markers and morphology, we performed two rounds of iterative indirect immunofluorescence imaging (4i)^44^ staining (round 1: cyclin A, cyclin B1, vimentin, DAPI, round 2: cyclin D1, DAPI) (Fig. 5a). We selected these proteins to determine the cell cycle phases: cyclin D1 as a marker for the G1 phase, cyclin A and cyclin B1 together as markers for the G2/M phase, and DAPI to determine DNA copy number and vimentin to identify EMT state. An inspection of the vimentin fluorescence intensity revealed that vimentin expression increased from day 0 to day 3, indicating that cells were beginning to undergo EMT (Fig. 5c, d). By day 14, there was a significant rise in vimentin intensity, suggesting that most cells had completed the EMT process (Fig. 5c, d).

**Figure 5.**
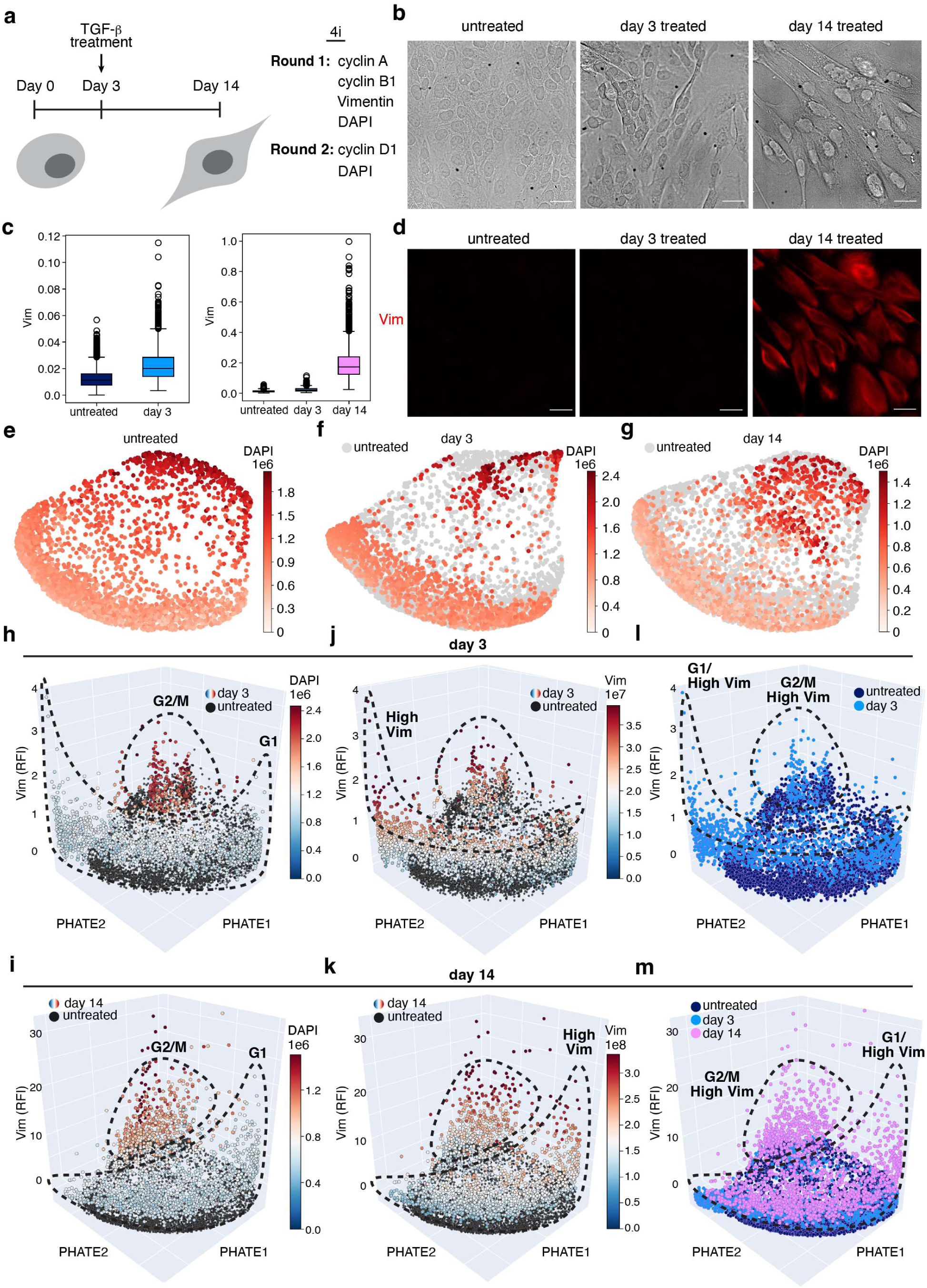
Protein expression reveals G1 and G2/M transition regions. (**a**) Experimental design: MCF10A cells were treated with TGF-β for 0, 3 and 14 days, followed by two rounds of 4i staining for cell cycle and EMT protein markers. (**b**) Representative differential interference contrast (DIC) images of MCF10A cells after 0, 3 and 14 days of TGF-β treatment. Scale bar, 30 μm. (**c**) Boxplots showing scaled vimentin expression (0 to 1) across treatment days. (**d**) Representative fluorescence images of vimentin expression in MCF10A cells after 0, 3 and 14 days of TGF-β treatment. (**e**-**g**) PHATE representation of untreated (**e**), day 3 (**f**), and day 14 (**g**) TGF-β treated MCF10A cells colored by DAPI intensity. Gray circles indicate untreated reference cells. (**h, i**) PHATE-vimentin representation of day 3 (**h**) and day 14 (**i**) treated cells colored by DAPI intensity. (**j, k**) PHATE-vimentin representation of day 3 (**j**) and day 14 (**k**) treated cells colored by vimentin. (**l, m**) PHATE-vimentin representation of day 3 (**l**) and day 14 (**m**) treated cells with the G1/high vimentin and G2/M-high vimentin regions labeled. The Vim z-axis indicates the relative fluorescence intensity (RFI).

We then learned a manifold representing cell proteomic states across the three time points with the features we extracted from 4i using PHATE to project the manifold into a 2D space for visualization. The PHATE representation revealed a clear circular cell cycle structure for the untreated cells (Fig. 5e, Fig. S8a). Using the untreated cell as a reference, we observed cells in day 3 and 14 in different stages of the cell cycle (Fig. 5f, g).

To examine the coupling between the cell cycle and EMT we utilized a 3D representation combining the 2D PHATE representation with vimentin intensity as the third axis. For day 3 and day 14, we observed two clusters corresponding to either the G1 or G2/M phase (Fig. 5h, i, Fig. S8b-j). Within each cluster, a subset of cells exhibited high vimentin intensity (Fig. 5j, k). The intersection between the G1 or G2/M groups with high vimentin intensity exhibited the coupling we computationally predicted previously between the cell cycle and EMT (Fig. 5l, m). In addition, cells with high vimentin intensity were not confined to a specific region within the G1 or G2/M phase but were broadly distributed, further supporting the loose coupling between the cell cycle and EMT. To view the interactive 3D plots, refer to Table S1.

In summary, multiplexed staining and manifold analysis confirmed our computational predictions from analyzing the scRNA-seq data. The proteomic data further corroborated that the cell cycle and EMT loosely couple during TGF-β induced EMT in MCF10A cells, resulting in cells broadly arresting from either the G1 or G2/M phase of the cell.

## Discussion

When a proliferating cell undergoes a cell phenotypic transition, it typically needs to temporarily pause or permanently exit the cell cycle to coordinate resource allocation^6^. Intuitively one expects that the exit takes place at only a few restricted regions, e.g., cell cycle checkpoints, of the cell cycle, while arrest can take place at various cell cycle stages. The present analysis of single-cell transcriptomic data combined with multiplex 4i protein staining data of MCF10A cells undergoing TGF-β induced EMT revealed that the arrest points were broadly distributed within the G1 or G2/M phases of the cell cycle.

The observed pattern can be visualized as a wheel and a lever attempting to stop the rotating wheel (Fig. 6). The rotating wheel represents the cell cycle, while the lever represents EMT. With increasing concentrations of TGF-β, more force is applied to the EMT lever eventually exerting pressure on to the wheel and disrupting cell cycle progression. When the EMT lever comes in contact with the rotating wheel, the cells in this region of the cell cycle halt their progression and transition from the epithelial to the mesenchymal state. This analogy illustrates how TGF-β treatment can induce widespread cell cycle arrest in either the G1 or G2/M phase^18–20^ while also promoting EMT. The difference in EMT programs between the two paths implies a potential mechanism for preferentially activating one path over the other, e.g., the G1 path at normal developmental and physiological conditions but not aberrant activation of the G2 path. The differential expression along the two paths may also contribute to heterogeneous cellular responses and drug resistance. Further studies are needed to elucidate the implications of the two EMT paths in the context of cell physiology and pharmaceutical intervention. Generally speaking, the observed coupling between cell cycle and EMT underscores the importance of quantitatively understanding the crosstalk between cell phenotypic transitions and the cell cycle when developing biomedical interventions.

**Figure 6.**
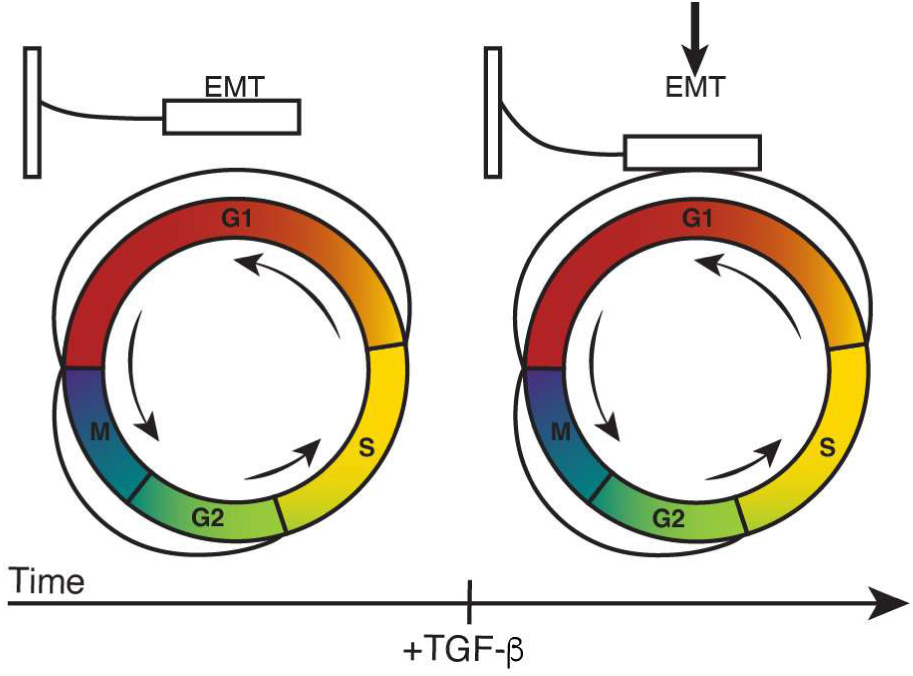
Schematic of the coupling between cell cycle and EMT consistent with observations in this study.

To perform single-cell data analysis, the standard procedure is to apply a dimensionality reduction method to high dimensional data to obtain a two-dimensional representation, for visualizing such complex data. Many dynamics inference methods such as trajectory inference are typically performed on low-dimensional representations of single-cell data, with an implicit assumption that the representation faithfully reflects the cellular processes being studied^45^. Single-cell omics approaches characterize cell state heterogeneity by quantifying the transcriptomic, proteomic, or epigenomic profiles of individual cells. Thus, a grand challenge is how to identify cellular processes relevant to a specific question of interest. A representation obtained with a *brute force* dimensionality reduction procedure, which emphasizes the largest contributions to heterogeneity and can conceal or distort the contributions of relevant processes, such as the cell cycle, as shown in this study. Furthermore, most dimensionality reduction methods do not include dynamic information, further worsening the situation. Our computational pipeline, which built upon the Revelio cell cycle framework^32^ to obtain the Cell Cycle Coordinate, underscores the importance of incorporating biological information when visualizing data in low-dimensional representations.

Examining cellular dynamics in a high dimensional space, aided by dynamical information such as a transcriptomic vector field, reduces mechanistic ambiguity. With the full dynamical model using RNA velocity information, we performed single-cell trajectory simulations in a high-dimensional state space and confirmed the existence of two distinct trajectories coupling cell cycle and EMT. Note that single-cell trajectories here differ from conventionally use of “trajectory” inference in the single-cell field. The latter typically refers to mean trajectories followed by an ensemble of cells that corresponds to the mean paths discussed in this work. In our case, single-cell trajectories refer to trajectories on a single-cell level.

This study demonstrates how biology-informed representations can reveal mechanistic insights into cellular processes from snapshot single-cell data, which are not transparent otherwise. Future studies can expand along several directions. We followed the Revelio model by assuming that one can approximate the single-cell data manifold to be cylindrical-shaped to describe cell cycle-dependent and -independent variations. However, this cylindrical-shape assumption is valid only locally. A data manifold with multiple cell types generally has more complex geometry which would require more sophisticated treatment.

In addition, we restricted our transition path analyses to a specific final mesenchymal state under several low concentrations of TGF-β. The MCF10A data at higher concentrations of TGF-β suggests the coexistence of different mesenchymal states, thus more than two types of EMT transitions paths may exist. Moreover, one can extend the trajectory analyses to investigate the coupling between various cellular processes such as EMT, gain or loss of stemness, and various other cellular fates such as apoptosis and senescence. Systematic identification of these different transition processes may require trajectory analyses without pre-selected start and end states. In short, improvements in cell state resolution and data sampling, combined with biology-informed in-silico trajectory analyses and experiments, enable systematic characterization and mechanistic insight into how different cellular processes couple to respond to one or multiple environmental and intracellular stimuli.

## Methods

### Processing of single-cell RNA-seq datasets

The MCF10A scRNA-seq dataset (GSE213753) consists of human mammary epithelial (MCF10A) cells that were treated with increasing concentrations of TGF-β (0, 12.5, 25, 50, 100, 200, 400, and 800 pM) for 14 days and then sequenced. The raw sequencing reads in the form of BAM files were first converted to fastq files using 10x Genomics Cell Ranger bamtofastq version 1.3.0 utility then Velocyto version 0.17.17 was used to obtain the unspliced and spliced count matrices.

### Single-cell RNA-seq analysis

After processing the MCF10A dataset, there was a total of 8983 cells. To obtain a shared UMAP representation for visual comparison between each of the TGF-β doses, all doses were normalized using *dynamo* version 1.0.0 “recipe_monocle” function where the n_top_genes = 1000, then PCA dimensionality reduction (n_components = 30) was performed, and finally UMAP (n_components = 2) was performed.

To calculate the EMT score we used a two-sample Kolmogorov-Smirnov (KS) test^34^ which compares the empirical cumulative distribution function (ECDF) of epithelial and mesenchymal gene sets using SciPy’s “ks_2samp” function. The KS test statistic and p-value for the ECDF_Epi_ ≥ ECDF_Mes_ and ECDF_Epi_ ≤ ECDF_Mes_ scenarios were calculated. A cell with a negative score is more epithelial while a cell with a positive score is more mesenchymal.

To determine which TGF-β doses contained epithelial cells for EMT analysis, we calculated the percentage of area overlap with the untreated data in the UMAP space. This was achieved by configuring an ellipse for each dose. First the mean coordinate was calculated to obtain the center of the ellipse. Then the eigenvalues from the covariance matrix were calculated to obtain the width and height of the ellipse. The angle of the ellipse was determined using the eigenvectors. To calculate the percentage of area overlap, the difference between the area of the untreated and treated data was taken and then divided by the area of the untreated data. The TGF-β doses with less than 5% area overlap were excluded from analysis.

For downstream analysis, each TGF-β dose was normalized separately “recipe_monocle” function where the n_top_genes = 2000 to calculate RNA velocity and then the corresponding vector field was obtained using *dynamo*. For downstream analysis, specifically for the 0, 12.5 and 25 pM TGF-β treated cells, a total of 3787 cells were analyzed.

To characterize EMT progression along the UMAP1 axis for both the 12.5 and 25 pM TGF-β treated cells, we performed Louvain clustering, resolution = 0.4, and obtained two clusters. Using scanpy’s version 1.9.8 function “rank_genes_groups” we obtained a ranking of the genes most representative of each cluster. With this list of genes, the gene expression of the top ranked genes for the 125 nearest neighbor cells to region a and region b was compared.

### Assigning cell cycle phase

Two methods were used to determine the cell cycle phase of each cell, *dynamo’s* “cell_cycle_score” function^28^ and Revelio’s “getCellCyclePhaseAssignInformation” function^32^. Both methods follow a similar approach, where each phase is defined by a list of genes. The average gene expression for each phase is calculated, z-normalized per phase and per cell, and the phase with the highest score is the assigned cell cycle phase. The difference between the two methods is that *dynamo’s* function assigns cells as either M, M/G1, G1/S, S or G2 while Revelio’s function assigns cells as either G1/S, S, G2, G2/M, or M/G1. These assigned discrete cell cycle phase was used to compare with the continuous cell cycle coordinate.

### Constructing the vector field from splicing-based RNA velocities

The python package, *dynamo* version 1.0.0, was utilized to obtain the vector fields^28^. *Dynamo* builds upon snapshot single cell data with RNA velocities inferred with various methods. For the datasets used in this study, we inferred the velocities from the unspliced (mRNA with introns) and spliced (mature mRNA without introns) RNA counts. The RNA turnover dynamics for a specific gene *i* can be described in the following ordinary differential equations^26^,

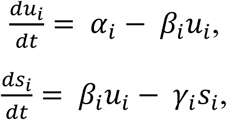

where *u_i_* represents the number of unspliced RNA transcripts for gene *i, s_i_* represents the number of spliced RNA transcripts of gene 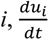 represents the change rate in unspliced RNA transcripts, 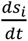 represents the change rate in spliced RNA transcripts, *α_i_* represents the transcription rate of gene *i, β_i_* represents the splicing rate constant of transcript *u_i_*, and *γ_i_* represents the degradation rate constant of transcript *s_i_*. Assuming a pseudo-steady state approximation with 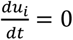, the conventional RNA velocity for a gene *i* can be defined as, 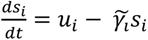, where 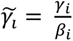 is a scaled gene specific degradation rate. RNA velocity can be estimated for each gene in each cell thus the state change rate of a cell can be represented by a high-dimensional vector of gene velocities. With a further assumption that all genes have the same splicing rate constant *β*, one can use the velocity vector to approximate the transition direction of individual cells in the transcriptomic state space.

### Projecting a vector field into a low-dimensional representation

For visualization purposes the high-dimensional velocities obtained from *dynamo* are often projected onto a two or three low-dimensional representation. This is commonly achieved by using Pearson, cosine kernels^26,27,46,47^, or mathematically rigorous GraphVelo method that is based on local linear embedding^47^ to observe cell fate transitions.

### Vector field analysis

*Dynamo’s* objective is to reconstruct a continuous vector field, i.e., a mechanistic systems biology virtual-cell model, from scRNA-seq gene expression and instantaneous RNA velocity data. The continuous vector field can be described as 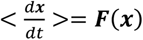, where ***x*** represents the gene expression state, 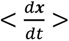 represents the RNA velocity at ***x*** averaged over its k-Nearest Neighbor (kNN) neighborhood, and ***F*** represents the learned vector field that maps the gene expression space to the velocity space, reflecting the gene-gene regulations of the system^28,40,47^.

With the continuous vector field, we can investigate the topology of the vector field through fixed point analysis. There are three types of fixed points: stable, unstable, and saddle points. Stable points are regions where the velocity vectors flow inward from all neighboring cells such as a stable cell type. Unstable points are regions where the velocity vectors only flow outward such as a pluripotent cell state which tends to differentiate quite quickly. Saddle points are regions where some velocity vectors flow inward and some outward such as a differentiated pluripotent cell which is relatively stable but also can further differentiate.

### Obtaining the one-dimensional Cell Cycle Coordinate

We used a finite temperature string method to define a one-dimensional cell cycle coordinate. The method was originally designed for defining a one-dimensional reaction coordinate that describes progression of a chemical reaction^41^ process, and has been adapted to analyze cell state transitions^16,40^. The finite-temperature string method assumes that all trajectories connecting a start and end region in the state space forms a narrow hyper-tube, and the center line of the hyper-tube is used to define a 1-D reaction coordinate that connects the initial and final regions through an iterative procedure.

While the finite-temperature string algorithm can be applied to find the 1-D path embedded in a multi-dimensional state space, here we restricted it to the DC1-DC2 plane defined by Revelio^32^. We designed an iterative procedure specifically for identifying the closed cell cycle coordinate (Fig. S3). Since the two DC modes contain most of the cell cycle information, the single cell data points already form a ring-shaped point cloud. We first calculated the center point of the data and generated a circle with a random radius from the center point as our initial guess of the cell cycle path. We then chose two consecutive points along this circle as our starting and ending points. Due to the iterative nature of this procedure the final path is not sensitive to this choice. Next, we iterated through the following steps,

1. We placed *N* points, including the starting and ending, called images, along the guessed cell cycle path uniformly with equal arclength distance. The image points divided the whole (2D) state space into Voronoi cells with each data point assigned to the Voronoi cell with the shortest Euclidean distance. For each Voronoi cell, the centroid of the single cell data was calculated and assigned as a corresponding new image point, except the starting and ending points.
2. With the new image points we then obtained a new guess of the continuous cell cycle path through cubic spline interpolation, and repeated step 1 until no more single cell data points were reassigned to Voronoi cells.
3. With the path from step 2, we chose an image point, typically far from the initial chosen starting and ending points, to calculate a new radius from the center of the data. With this new radius a circle was generated, as an improved initial guess, and two consecutive points were selected as the starting and ending points. With this improved guess the above two steps are repeated. One can repeat such re-optimization, but in practice we found one round was sufficient.

With the final image path from step 3, each cell was assigned a Cell Cycle Coordinate value through interpolation between two nearest image points.

### Determining the new cell cycle representation

To obtain the final two-dimensional representation that allowed us to observe how the cell cycle couples with other biological processes, a second axis representing the biological process of interest was needed. The cell by DC matrix output by Revelio showed that the first two dimensions capture a majority of the cell cycle dependent variation therefore the remaining dimensions must contain the cell cycle independent variation, most likely the biological process being studied. The third DC dimension was designated as the new y-axis to create the new two-dimensional representation for the untreated, 12.5 and 25 pM TGF-β treated cells.

### Generating gene expression heatmap and expression plots along the Cell Cycle Coordinate

Since we identified two regions, a quiescent and cycling region, in the untreated MCF10A cells, we selected for only cycling cells to accurately capture the gene expression of cell cycle related genes along the Cell Cycle Coordinate. To generate the heatmap the gene expression of several G1 and G2 genes were averaged along each point of the Cell Cycle Coordinate. Cells with zero expression were dropped and each gene was normalized using standard scaling. Similarly for the expression plots, for each gene the expression was averaged along each point of the Cell Cycle Coordinate, with zero expression cells dropped. The smoothed dashed line representing a genes expression pattern used SciPy’s “make_smoothing_spline” function with the regularization parameter set to 1000.

### Performing trajectory simulations

The trajectory simulations utilize a transition matrix defined in 30-d PCA space. The transition matrix was obtained from *dynamo* using the cosine kernel method^26,28^. The method uses RNA velocity information to estimate the transition probability (*p_ij_*) from a cell state *i* to another neighboring state *j* in the 30-d state space, 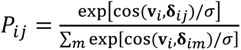, with **v***_i_* the RNA velocity vector of cell state *i*, **δ***_ij_* = ***x****_j_* − ***x****_i_* where ***x****_i_* and ***x****_j_* the expression states of cell state *i* and *j*, cos( , ) denoting the cosine similarity between two input vectors, *σ* an arbitrary bandwidth parameter (assumed a value of 10 in this study), and the sum is over all kNN neighbors of cell *i*.

Following chemical rate theory, we performed transition path theory analysis^16,40,48^ as the following. Aided with the 2D CC-EMT coordinate representation, we chose two threshold values of the EMT coordinate (*y*), *∈*_1_ < *∈*_2_, d divided the state space into three regions, Epithelial (*y* < *∈*_1_), Intermediate (*∈*_1_ ≤ *y* ≤ *∈*_2_), and Mesenchymal ((*y* > *∈*_2_),). We varied the values of *∈*_1_ and *∈*_2_, and found the subsequent analyses were insensitive to the choice, a feature common in transition path theory analysis^48^. Using the transition matrix, we randomly selected experimentally sampled single cell data within the Epithelial region as starting points and launched Gillespie trajectory simulations^49^ in the full-dimensional state space. Following transition path theory, we termed trajectories starting from the Epithelial region and ending at the Mesenchymal region without returning to the Epithelial region as reactive trajectories. Trajectories that returned to the start region before hitting the end region, termed non-reactive trajectories, were discarded.

### Clustering and identifying paths in high dimensional space

For both the 12.5 pM and 25 pM TGF-β treated MCF10A cells, ten rounds of 100 reactive trajectories were acquired, totaling 1000 trajectories for each dose. For each trajectory simulation round, the “TimeSeriesKMeans” (n_clusters, metric=softdtw, max_iter=10, random_state=0, init=k-means++) function from tslearn version 0.6.3 was applied to cluster the trajectories in the 10-dimensional DC space. The number of clusters, n_clusters, was user defined, but we checked for two and three clusters to ensure that there were only two clusters.

To obtain a mean trajectory for each cluster, we used an adapted finite temperature string method from Wang et al^16,40^. The procedure was similar to what was used in obtaining the Cell Cycle Coordinate, but with two major differences. First, the procedure was performed in the leading 10-d DC space. Second, the reactive trajectories also have weights on determining the image points in each Voronoi cells, as detailed in^16,40^.

To map the mean trajectories from DC space to other spaces (e.g., CC-EMT) the nearest neighbor cell in DC space was identified and its corresponding coordinate in the target space was used. To compare the mean trajectories obtained in 10-d DC space from each round of trajectory simulations, fastdtw version 0.3.4 was utilized to calculate the Euclidean distance between two trajectories. Pairwise comparisons of all mean trajectories generated a heatmap revealing the number of clusters.

### Differential gene expression analysis of the two paths

With one set of mean trajectories from one round of 12.5 pM TGF-β trajectory simulations more points were interpolated using (Fig. S4d) SciPy’s UnivariateSpline function, with a smoothing factor of 0.5, obtaining x coordinates given a set of coordinates along the EMT axis. Having a shared EMT axis allows us to make a one to one comparison between the two paths. With these interpolated mean trajectories for the G1 and G2/M paths in CC-EMT space we identified the corresponding cell states or nearest neighbor in DC space and then converted it into gene expression space using a rotation matrix obtained from Revelio^32^.

### Cell Culture

Human mammary epithelial cells (MCF10A; ATCC, CRL-10317) were cultured in DMEM/F-12 medium supplemented with 5% horse serum, 20 ng/mL epidermal growth factor (EGF), 0.01 mg/mL insulin, 500 ng/mL hydrocortisone, and 100 ng/mL cholera toxin. Cells were maintained in a humidified atmosphere of 5% CO₂ at 37°C and utilized within 10 passages.

For the TGF-β treatment, 1×10^5^ MCF10A cells were plated in 35mm glass-bottom plates (Ibidi, #80137) and cultured for two to three days. When cells reached 70-80% confluency, 0.55 ng/mL TGF-β was added to the culture media. The culture media with TGF-β was refreshed every 24 hours. TGF-β treated cells were fixed with 4% PFA after 3 and 14 days respectively. The fixed cells were then stored at 4 °C for immunofluorescence staining.

### 4i staining and imaging

Iterative Indirect Immunofluorescence Imaging (4i) staining was performed following the protocol published by Kramer et. al^50^. The immunofluorescence staining was carried out in two rounds: round 1: cyclin A (1:50, Santa Cruz, sc-271682), cyclin B1 (1:100, R&D, AF6000), vimentin (1:150, Thermofisher, PA5-27231), and DAPI; round 2: cyclin D1 (1:100, Santa Cruz, sc-20044) and DAPI. A Nikon Ti-E2 microscope was used to image under 40X (0.95NA). The following channels were used: DAPI, TRITC, FITC, Cy5. The final image was stitched from 8×8 tiles (each 1024×1024 pixels) with a stitching overlap of 15% to get one 7117×7117-pixel image. All imaging for each channel staining marker was conducted using identical microscope settings and performed on the same day.

### Image analysis

For each image, we first subdivided each 7,117×7,117-pixel field into a 4×4 grid of 16 tiles as input to the pre-trained Cellpose cyto3 model (3.0.9.dev8+gc03958c)^51^. Segmentation was performed on each tile using Differential Interference Contrast (DIC) and vimentin channels to generate cell masks. The DAPI channels were used to generate nuclear masks. Nuclear masks required no manual correction due to the high signal-to-noise ratio of DAPI, however cell contours were manually refined using the DIC channel as reference. Subtle offsets between the two rounds of imaging were corrected by registering nuclear masks with a phase cross correlation algorithm in the scikit-image package version 0.20.0. We then applied the same shift parameters to the other channels. Channel-specific background intensities were measured in ImageJ and subtracted from every image.

Multiple features were extracted from the images, including the ratio of nuclear mean intensity over cytoplasmic mean intensity for cyclin D1, A, and B1, total nuclear DAPI and total cell vimentin intensity. All features except the total cell vimentin intensity from the three independent experiments were pooled and z-score normalized. Using these features, we applied PHATE version 1.0.11 (k-nearest neighbors = 45) to learn a two-dimensional shared manifold representing cell states across all three time points. Total cell vimentin intensity was then overlaid as the third axis to visualize EMT progression across different days of treatment in a 3D representation.

Clustering was run separately for the three groups: no treatment, 3, and 14 days TGF-β treated. We used scikit learn’s KMeans clustering with k set to 2 and all other parameters kept at their default settings. The input to KMeans was a one column array containing the DAPI sums for all cells in the given group. After fitting, the two clusters were ordered based on their mean DAPI intensity to maintain a consistent labeling across groups.

## Data availability

All the raw sequencing data for the MCF10A dataset is available in the NCBI Gene Expression Omnibus under the accession <u>GSE213753</u>. The features extracted from 4i can be found in the supplemental.

## Code availability

All code can be found on GitHub https://github.com/xing-lab-pitt/two-paths.

## Acknowledgements

The study was supported by NIDDK R01 DK119232 (JX), R56 DK (JX), NIGMS R01 (JX), NSF 2325149 (JX), NSF 2205148 (JX) and the pilot award from the University of Pittsburgh Hillman Cancer Center (to JX). We would like to thank Weikang Wang for the helpful discussions and Kazuhide Watanabe for providing the data.

## Author contributions

Author contributions were determined using the CRediT model: Conceptualization: SH, JX. Formal analysis: SH, GYu, ZZ, KN. Funding acquisition: JX. Investigation: SH, YL, GYu, ZZ, KN. Methodology: SH, YL, GYu, ZZ, JZ, YZ, JX. Project administration: JX. Supervision: JX. Visualization: SH, GYu, ZZ. Writing – original draft: SH, JX. Writing – review & editing: SH, YL, GYu, ZZ, AG, GYao, JX.

## Box 1.

**Dimensionality reduction may distort the underlying dynamics**

For illustrative purposes, consider that the cellular dynamics of proliferative cells is described by a 3D vector field on a hollow cylindrical manifold, where the circle of the cylinder depicts the cell cycle and the straight side depicts additional cellular processes (Box Fig. 1a). Such a geometric manifold is topologically impossible to be faithfully represented in a two-dimensional space without slicing or folding the manifold onto a flat surface. This is exemplified when one applies various dimensionality reduction methods, e.g., Uniform Manifold Approximation and Projection (UMAP)^35^, Principal Component Analysis (PCA), diffusion Map^52^, and Potential of Heat-diffusion for Affinity-based Trajectory Embedding (PHATE)^53^. We projected the vector field into 2D using a cosine kernel^26^, a method commonly used to project a vector field to a lower dimensional representation. The resulting 2D UMAP representations heavily distorted the manifold (Box Fig. 1b). Vectors parallel in the original 3D space appeared as converging into some spurious fixed points in the UMAP space. In the case of PCA or diffusion map, the resulting 2D representations were only able to capture one side of the cylinder (Box Fig. 1c, d). PHATE, notably, was able to capture the circle in the manifold and partially capture the depth of the cylinder (Box Fig. 1e). One plausible reason for the best performance of PHATE is that the algorithm considers both local and global structures and utilizes Multidimensional scaling (MDS) to ensure the best visualization in 2D. In comparison, PCA emphasizes the global structure, UMAP has issues with distances with no inherent meaning, and diffusion map takes local and global structure into account but is not optimal for 2D visualization.

In this simple case, we have the ground truth vector field as reference to compare to the projected vector field. An actual scRNA-seq data manifold likely has a more complex structure, which imposes challenges for detecting distortion of the manifold due to folding and compression that arise during projection to a lower-dimensional representation. These distortions obscure underlying biological processes and skew the velocity vector field. Incorporating biological context provides additional clues for proper projection, however ultimately analysis of dynamics with the original representation is still needed.

**Box Fig. 1.**
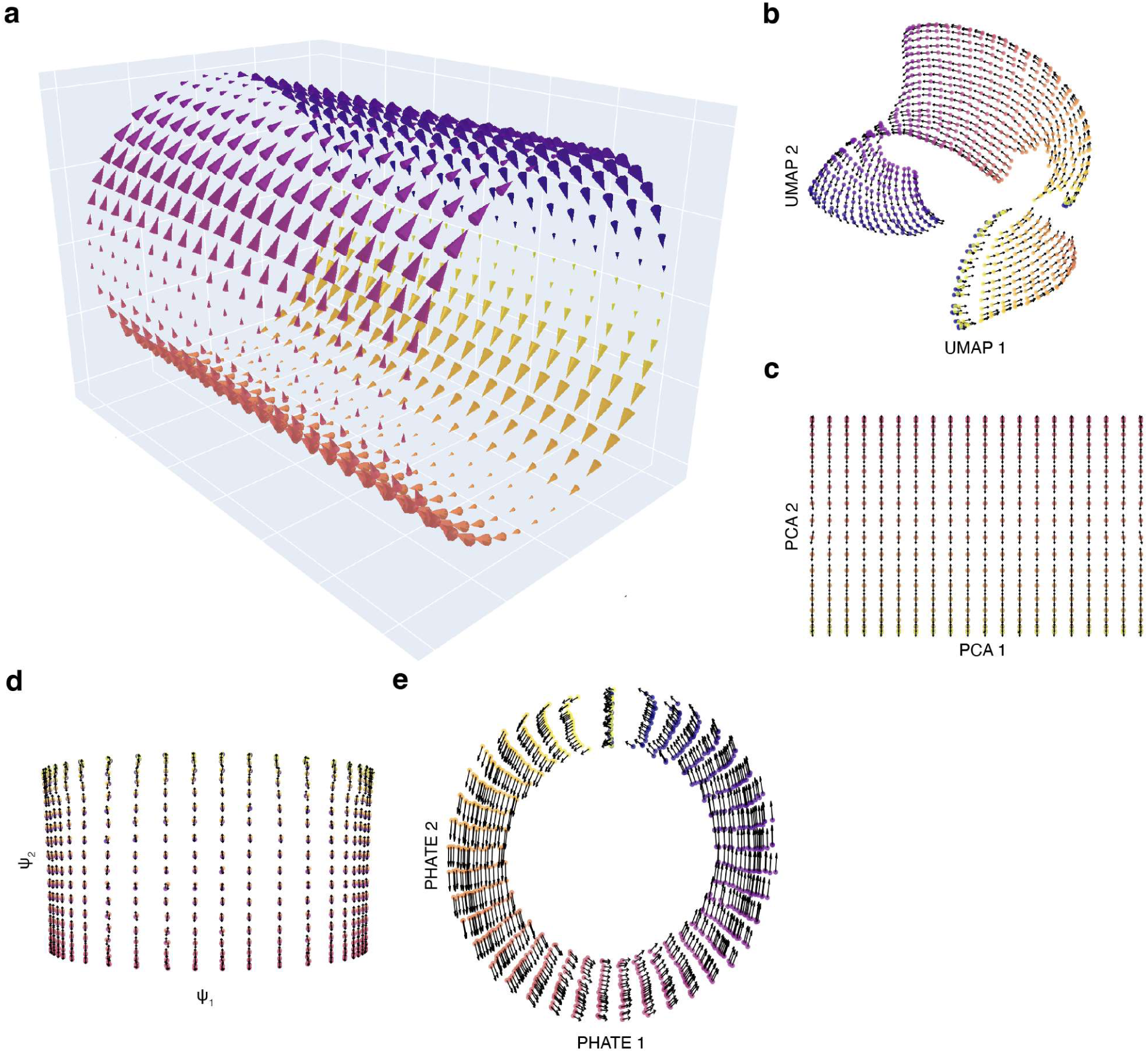
Applying different dimensionality reduction methods to a 3D vector field. (**a**) 3D cylindrical vector field. The 3D vector field projected to (**b**) 2D UMAP space, (**c**) 2D PCA space, (**d**) 2D diffusional map space and (**e**) 2D PHATE space using cosine kernel.

## Supplemental Figures

**Supplemental Figure 1.**
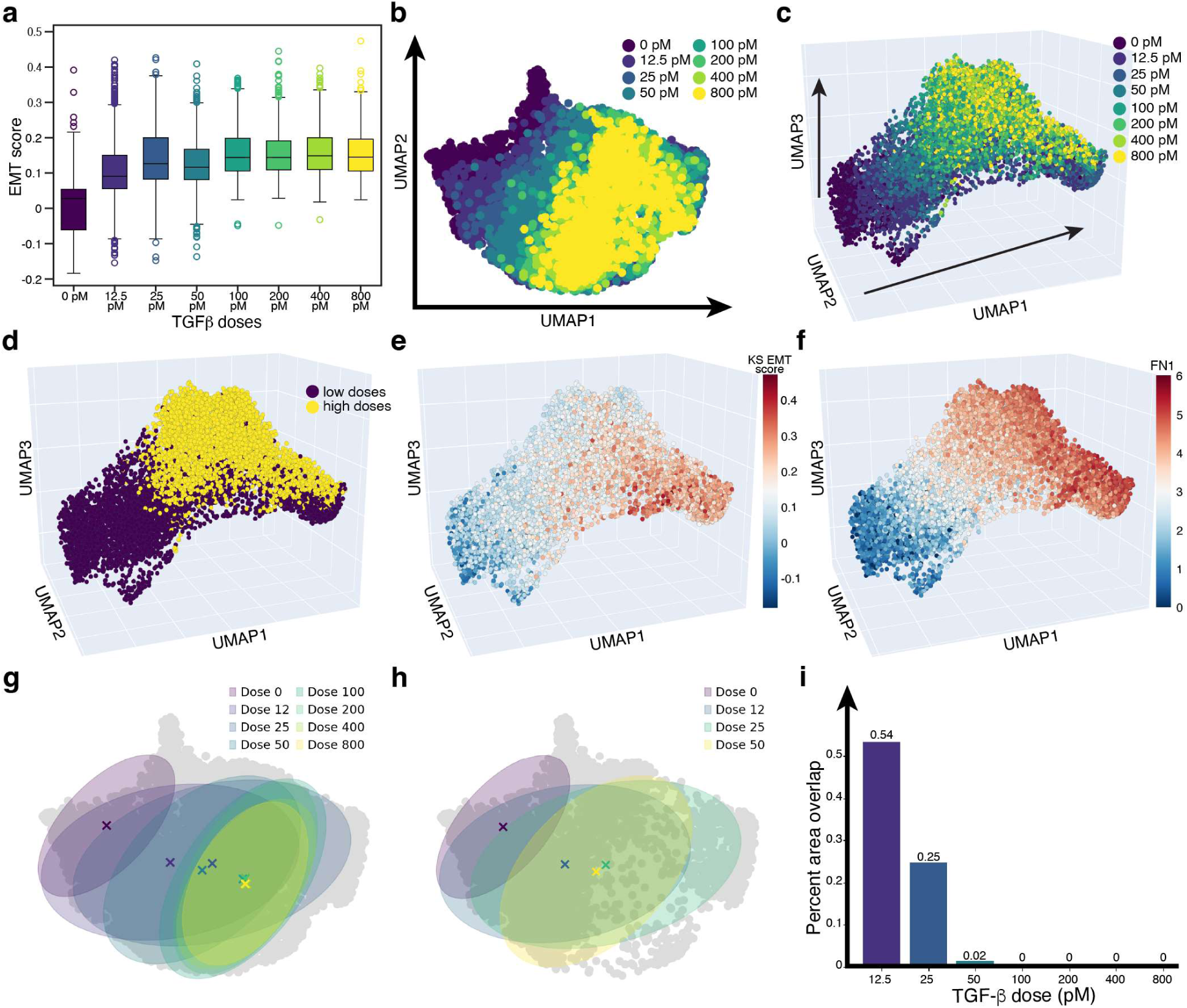
EMT score analyses of MCF10A scRNA-seq dataset. (**a**) KS EMT score for each TGF-β dose. (**b**-**c**) All TGF-β doses (0 pM, 12.5 pM, 25 pM, 50 pM, 100 pM, 200 pM, 400 pM, 800 pM) in the two-dimensional and three-dimensional UMAP space colored by TGF-β dose. (**d**) scRNA-seq data colored by TGF-β dose. Low doses: 0, 12.5, 25 pM TGF-β. High doses: 50, 100, 200, 400, 800 pM TGF-β. (**e-f**) scRNA-seq data colored by the KS EMT score (**e**) (lower KS score = more epithelial and higher KS score = more mesenchymal) and FN1 gene expression (**f**). (**g**-**i**) The percent area overlap between each TGF-β dose and the untreated MCF10A condition.

**Supplementary Figure 2.**
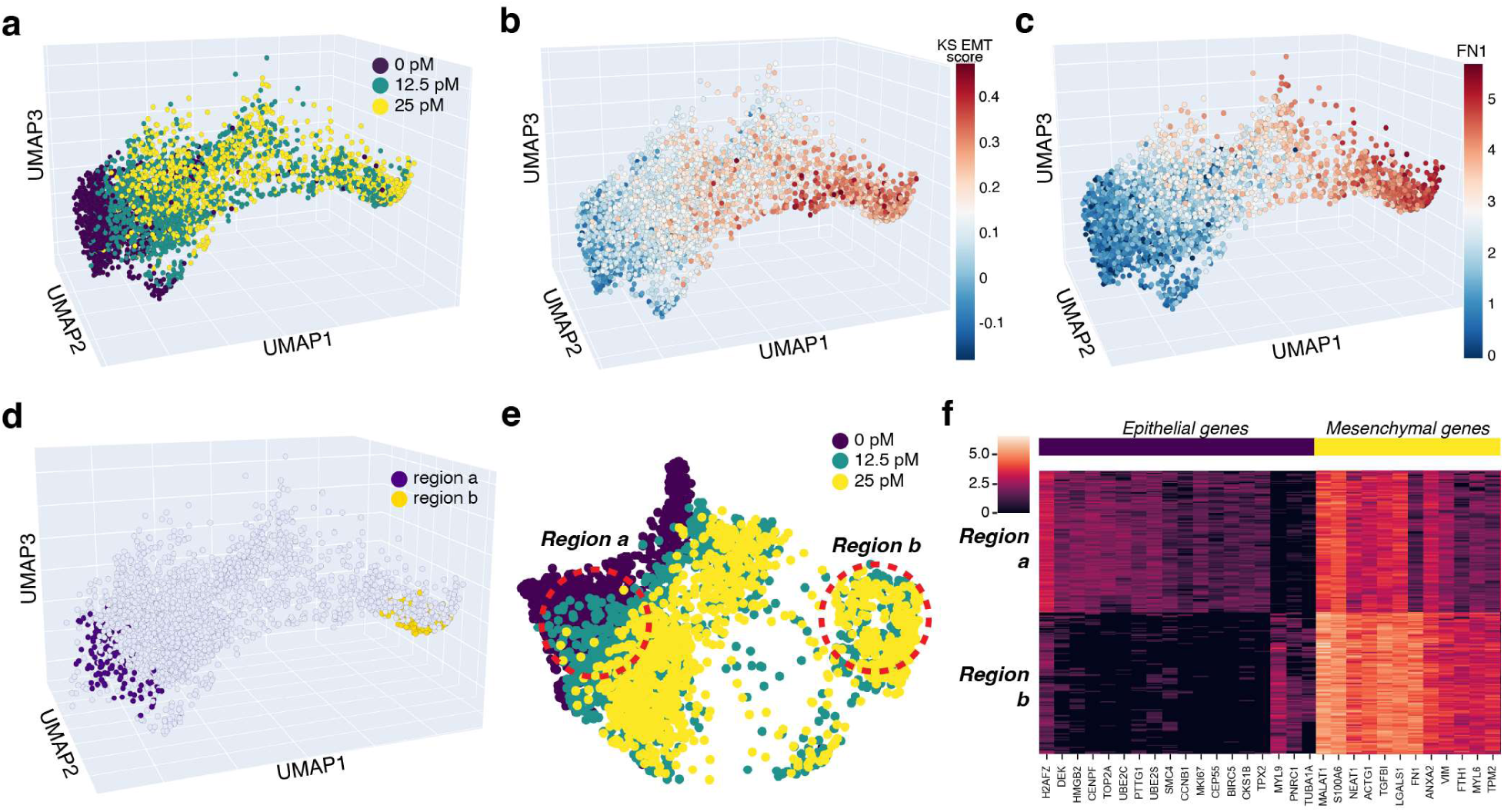
EMT score analyses of MCF10A cells treated with low doses of TGF-β. (**a**) scRNA-seq data colored by TGF-β dose. (**b**) scRNA-seq data colored by KS EMT score (lower KS score = more epithelial and higher KS score = more mesenchymal) and (**c**) FN1 gene expression. (**d**) 3D and (**e**) 2d UMAP representation of 0, 12.5, 25 pM TGF-β treated cells with two highlighted regions utilized for differential gene expression analysis. (**f**) Gene expression comparison between region a and region b looking at epithelial and mesenchymal marker genes.

**Supplemental Figure 3.**
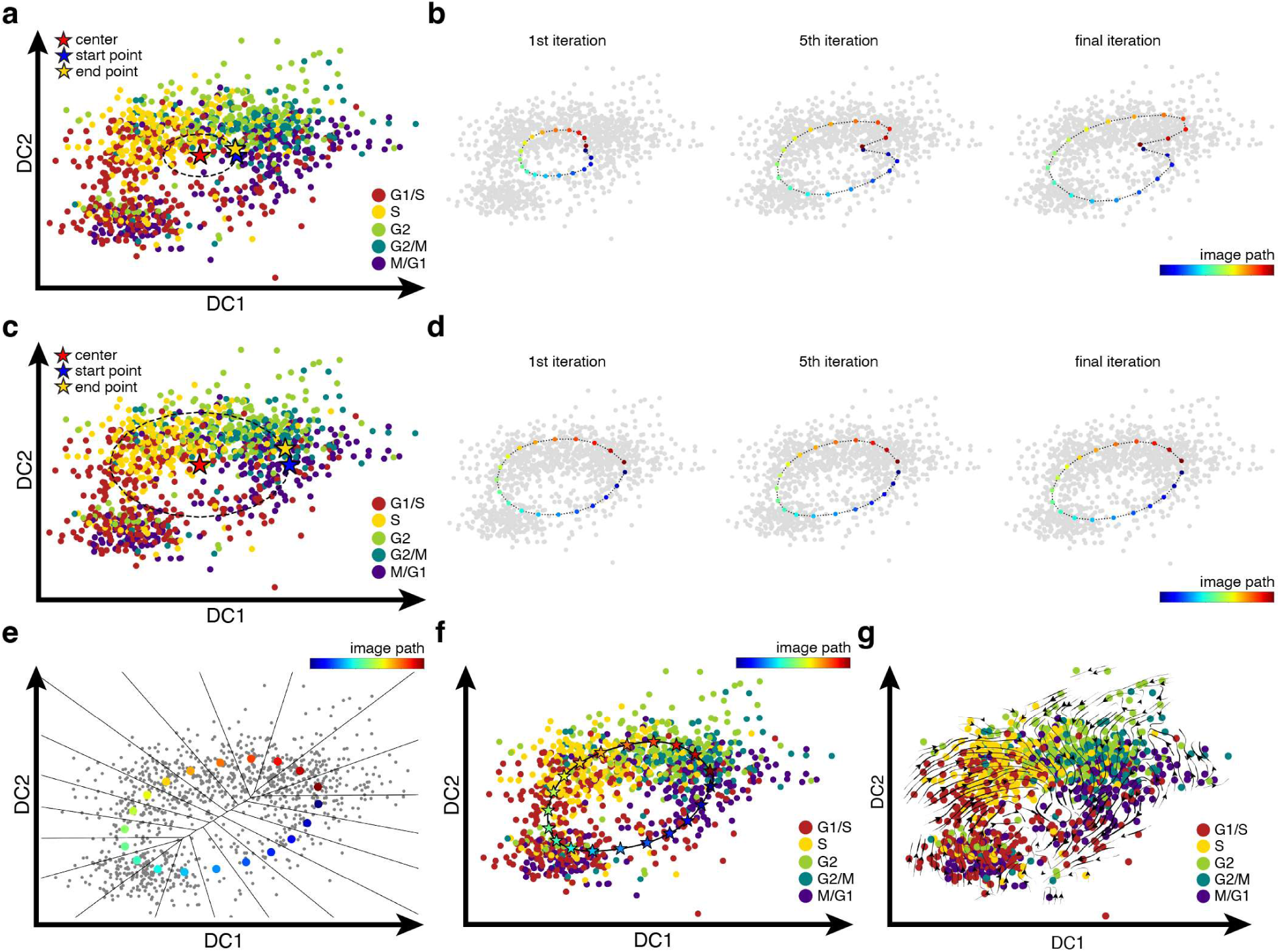
Finite temperature string method for identifying the Cell Cycle Coordinate. (**a**) Untreated MCF10A cells in DC space. The dashed line indicates the initial circle guess centered on the data. The starting and ending points correspond to two consecutive points along the circle. (**b)** Several iterations of the image path obtained during the application of the finite temperature string method. (**c**) A more refined circular guess is constructed using the last iteration from the first application of the finite temperature string method. (**d**) Several iterations of the image path obtained during the second application of the finite temperature string method. (**e**) The final image path divides the DC space into Voronoi cells. (**f**) With the final image path, points in between images are interpolated to obtain the Cell Cycle Coordinate. (**g**) Vector field obtained with the untreated MCF10A cells in DC space. Cell cycle phases were assigned using Revelio.

**Supplemental Figure 4.**
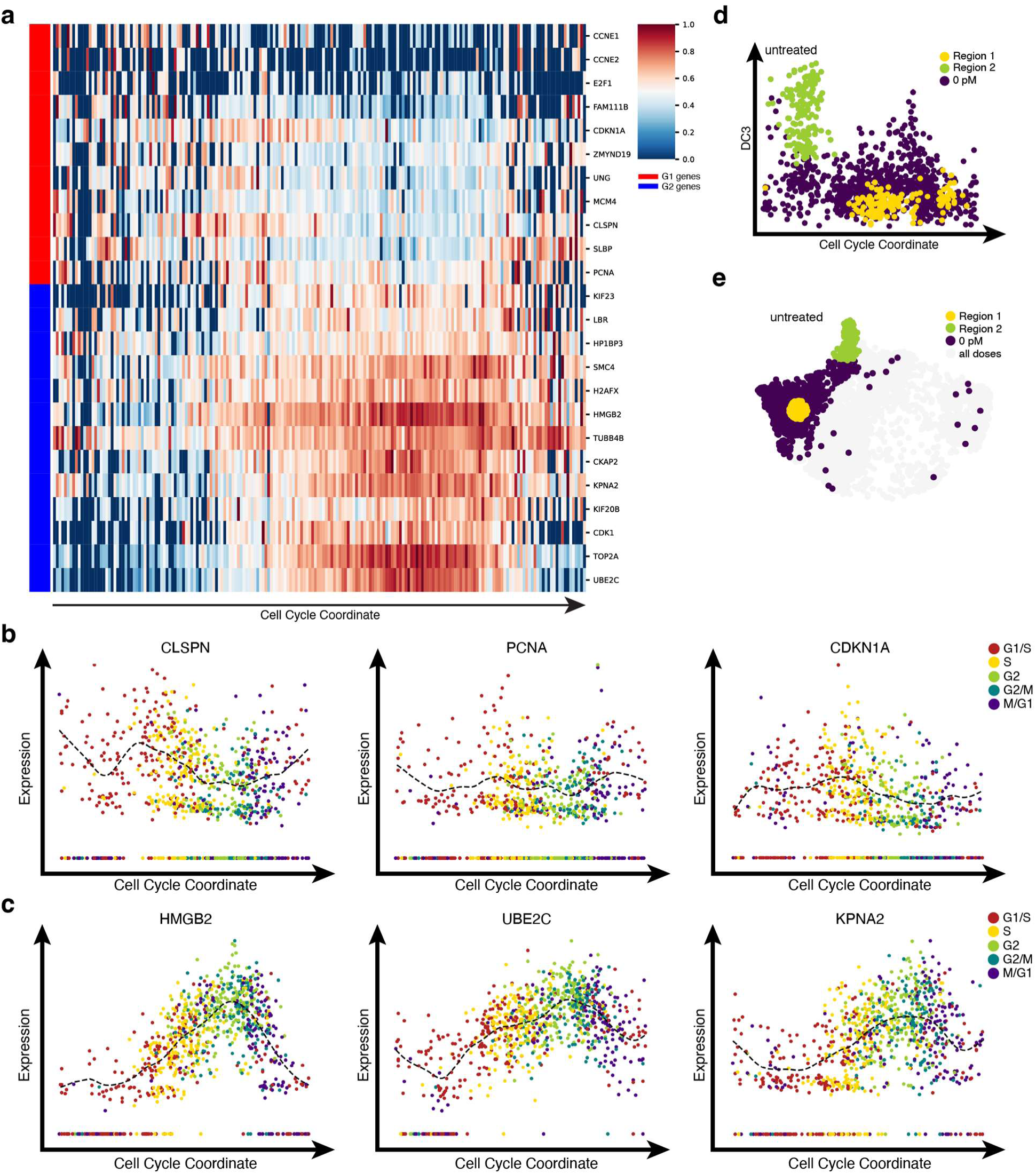
Cell cycle phase profile along the Cell Cycle Coordinate in untreated MCF10A cells. Only cycling cells were selected for this analysis. (**a**) Heatmap of G1 and G2 gene expression along the Cell Cycle Coordinate. The expression for each gene was normalized from 0 to 1. (**b**) Expression plots of G1 genes CLSPN, PCNA, and CDKN1A. (**c**) Expression plots of G2 genes HMGB2, UBE2C, and KPNA2. Cell cycle phases were assigned using Revelio. The dashed lines were produced using B-splines to smooth out the expression pattern. (**d**) Single cell data points in the Cell Cycle Coordinate-EMT representation. The colors indicate the cells chosen for differential gene expression analysis. (**e**) Single cell data points in the Cell Cycle Coordinate-EMT space colored by the corresponding quiescent and cycling regions. Regions 1 and 2 in panels d and e correspond to the regions shown in Fig. 1d.

**Supplemental Figure 5.**
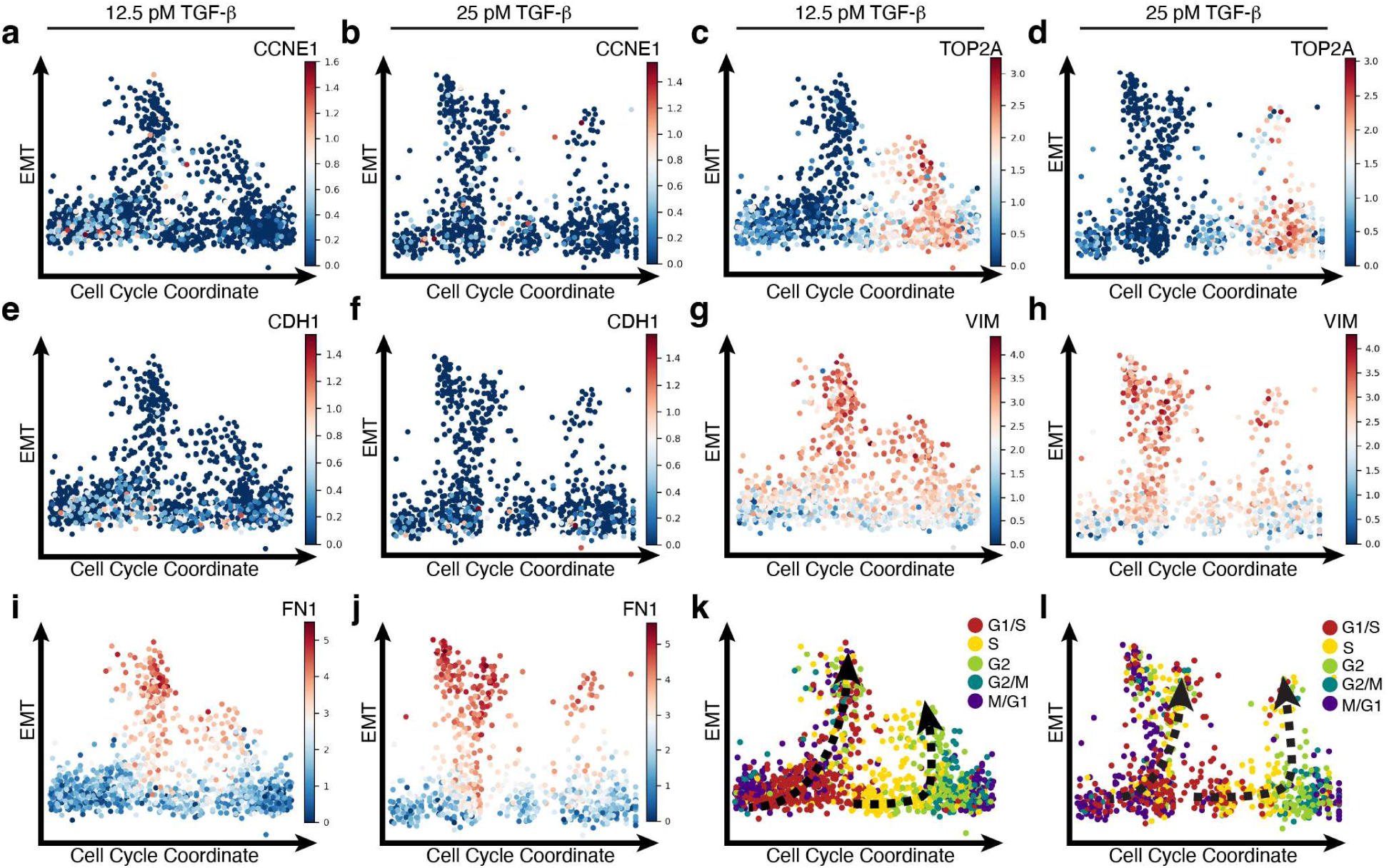
Cell Cycle Coordinate-EMT visualization of selected genes and cell cycle phase in scRNA-seq data of MCF10A cells treated with 12.5 and 25 pM TGF-β. (**a**, **b**) CCNE1; (**c**, **d**) TOP2A; (**e**, **f**) CDH1; (**g**, **h**) VIM; (**i**, **j**) FN1; (**k**, **l**) cell cycle phases were assigned using Revelio. The dashed lines indicate the two potential paths.

**Supplemental Figure 6.**
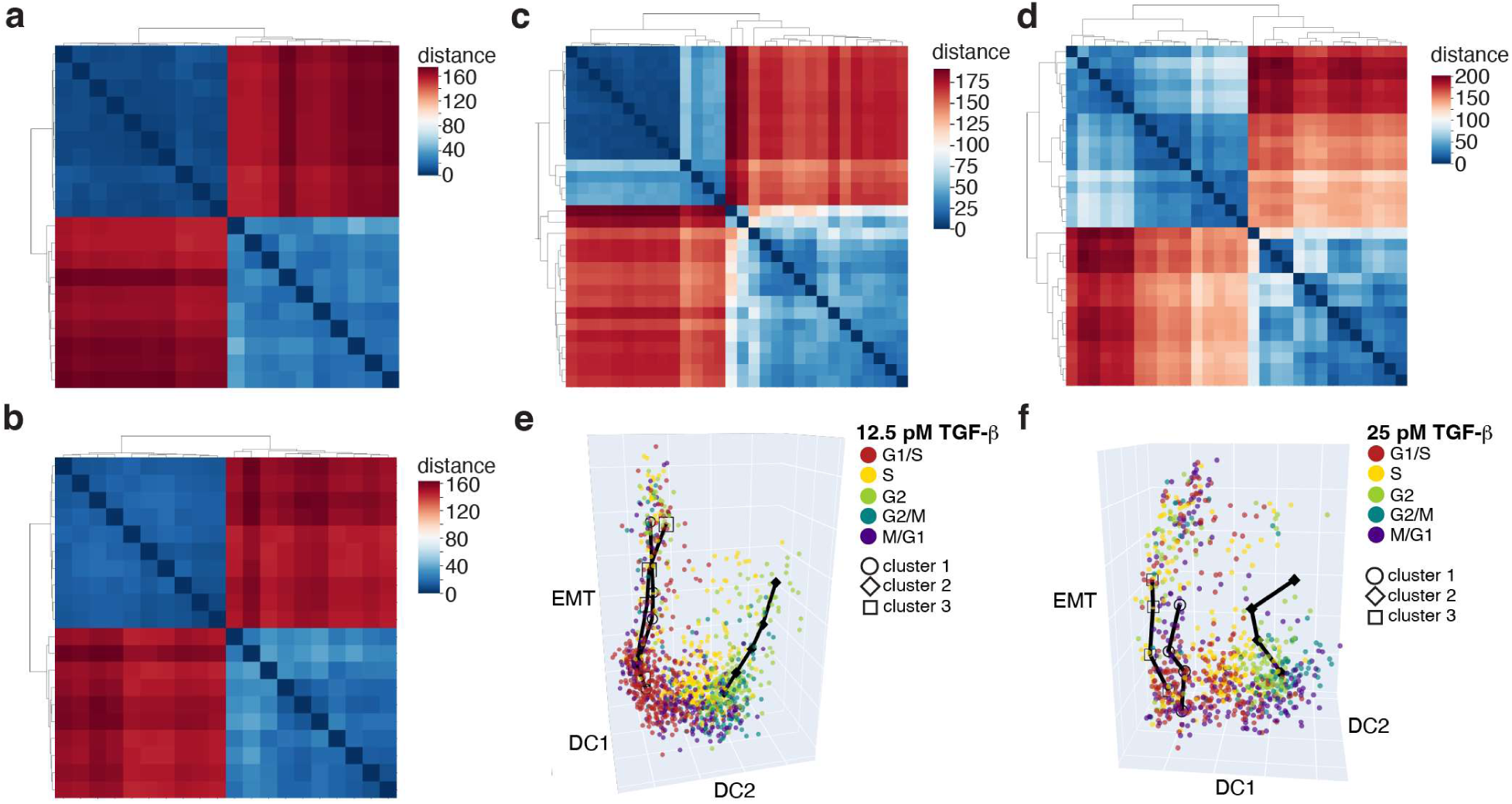
Clustering of simulated EMT trajectories. **(a,b)** Distance matrices of the 20 mean trajectories generated from trajectory simulations showing two clusters for (**a**) 12.5 pM and (**b**) 25 pM TGF-β. Distance matrices of the 30 mean trajectories generated from trajectory simulations for (**c**) 12.5 and (**d**) 25 pM. The three representative trajectories from one (**e**) 12.5 pM and (**f**) 25 pM trajectory simulation in the DC-EMT space. Cell cycle phases were assigned using Revelio.

**Supplemental Figure 7.**
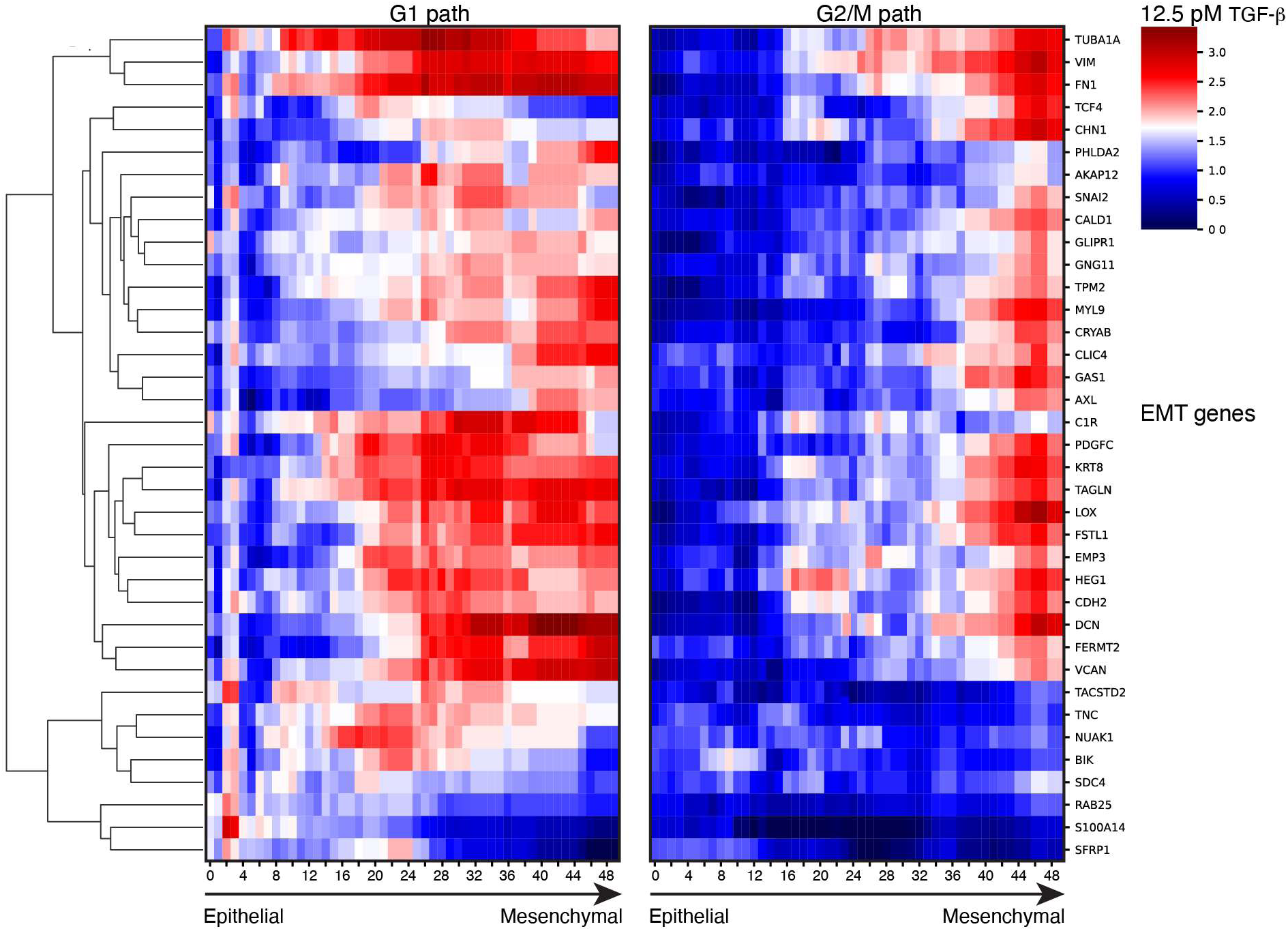
Differential gene expression analysis of EMT genes between the two paths in 12.5 pM TGF-β treated cells. The x-axis represents the same EMT axis with increased EMT progression from left to right.

**Supplemental Figure 8.**
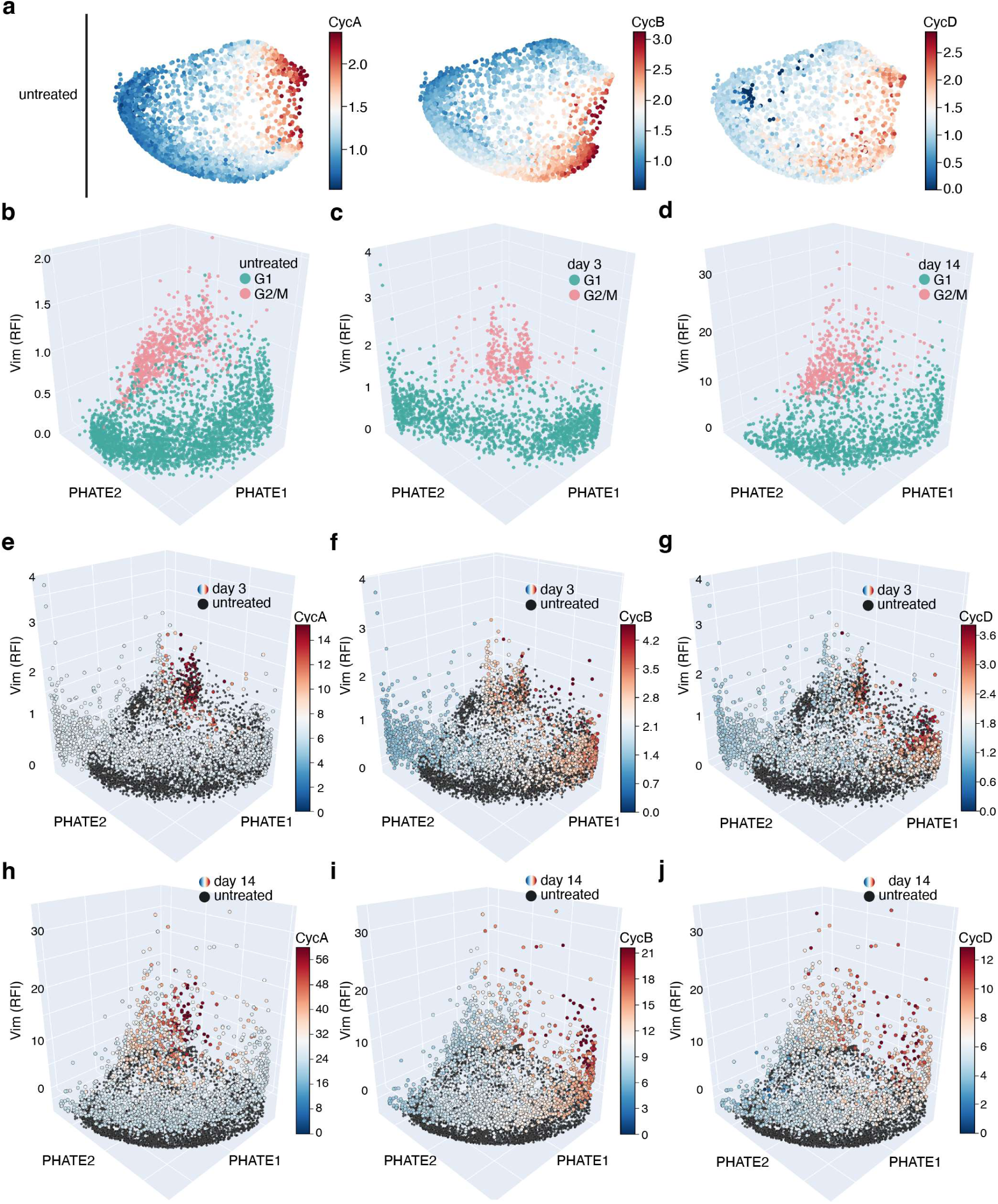
Additional analysis of MCF10A 4i data. (**a**) PHATE visualization of cyclin A, cyclin B1, and cyclin D1 protein expression in untreated MCF10A cells. (**b**-**d**) K-means clustering identified two groups corresponding to the G1 and G2/M phase for the untreated, day 3, and day 14 TGF-β treated cells. (**e**-**f**) 3-d visualization of (**e**) cyclin A, (**f**) cyclin B1, and (**g**) cyclin D1 protein expression in MCF10A cells treated with TGF-β treatment for three days. (**h**-**j**) 3-d visualization of (**h**) cyclin A, (**i**) cyclin B1, and (**j**) cyclin D1 protein expression in MCF10A cells treated with TGF-β treatment for 14 days. The black dots are the untreated cells as reference. The Vim z-axis indicates the relative fluorescence intensity (RFI).

**Table S1.**
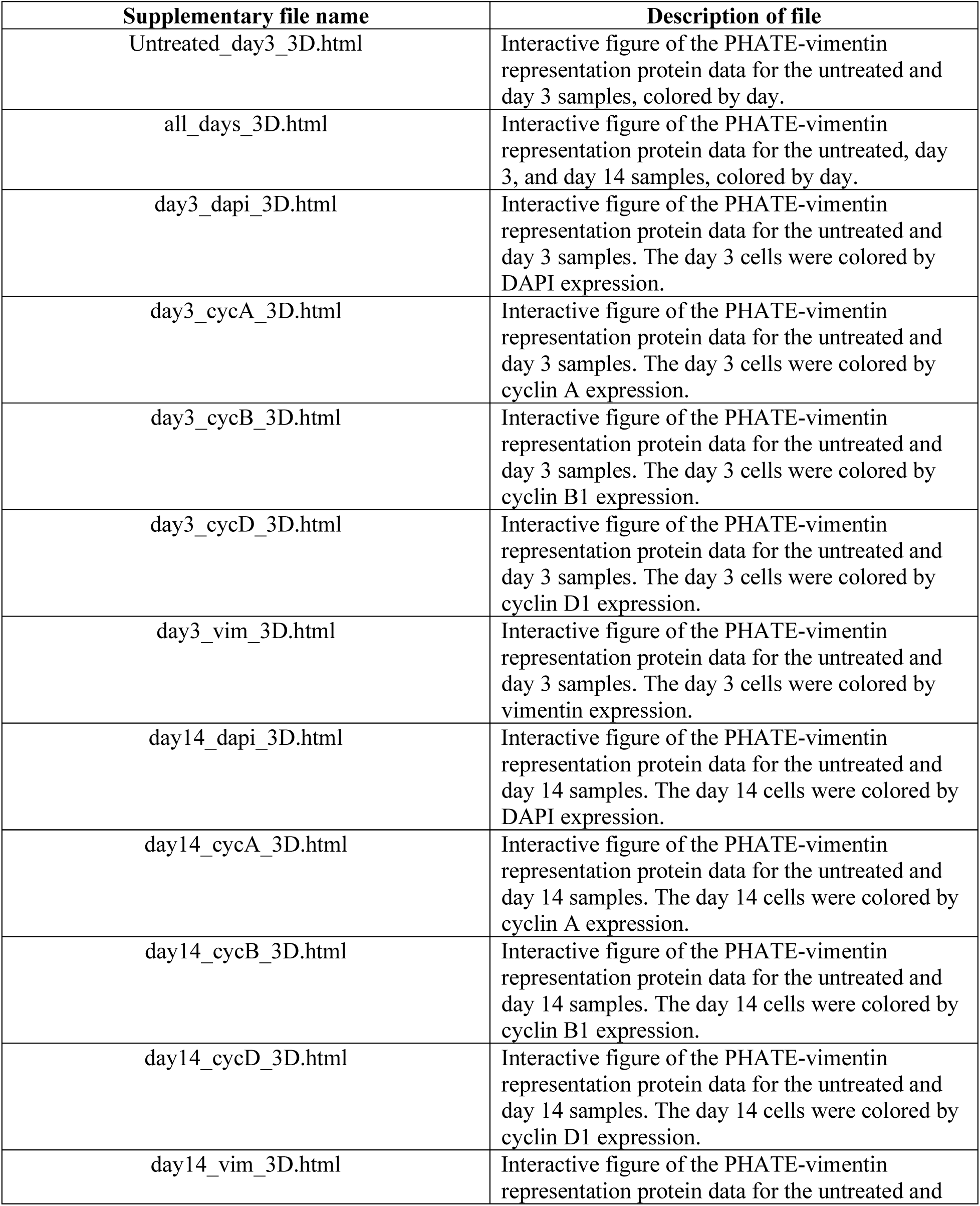
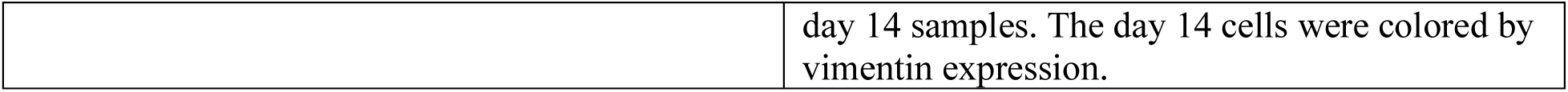
List of interactive plots related to 4i analysis.

| Supplementary file name | Description of file |
| --- | --- |
| Untreated_day3_3D.html | Interactive figure of the PHATE-vimentin representation protein data for the untreated and day 3 samples, colored by day. |
| all_days_3D.html | Interactive figure of the PHATE-vimentin representation protein data for the untreated, day 3, and day 14 samples, colored by day. |
| day3_dapi_3D.html | Interactive figure of the PHATE-vimentin representation protein data for the untreated and day 3 samples. The day 3 cells were colored by DAPI expression. |
| day3_cycA_3D.html | Interactive figure of the PHATE-vimentin representation protein data for the untreated and day 3 samples. The day 3 cells were colored by cyclin A expression. |
| day3_cycB_3D.html | Interactive figure of the PHATE-vimentin representation protein data for the untreated and day 3 samples. The day 3 cells were colored by cyclin B1 expression. |
| day3_cycD_3D.html | Interactive figure of the PHATE-vimentin representation protein data for the untreated and day 3 samples. The day 3 cells were colored by cyclin D1 expression. |
| day3_vim_3D.html | Interactive figure of the PHATE-vimentin representation protein data for the untreated and day 3 samples. The day 3 cells were colored by vimentin expression. |
| day14_dapi_3D.html | Interactive figure of the PHATE-vimentin representation protein data for the untreated and day 14 samples. The day 14 cells were colored by DAPI expression. |
| day14_cycA_3D.html | Interactive figure of the PHATE-vimentin representation protein data for the untreated and day 14 samples. The day 14 cells were colored by cyclin A expression. |
| day14_cycB_3D.html | Interactive figure of the PHATE-vimentin representation protein data for the untreated and day 14 samples. The day 14 cells were colored by cyclin B1 expression. |
| day14_cycD_3D.html | Interactive figure of the PHATE-vimentin representation protein data for the untreated and day 14 samples. The day 14 cells were colored by cyclin D1 expression. |
| day14_vim_3D.html | Interactive figure of the PHATE-vimentin representation protein data for the untreated and |

**Table S1. List of interactive plots related to 4i analysis.**
|  |
| --- |
| day 14 samples. The day 14 cells were colored by vimentin expression. |

